# Nudging the N170 forward with prior stimulation – Bridging the gap between N170 and recognition potential

**DOI:** 10.1101/2020.10.06.327908

**Authors:** Canhuang Luo, Wei Chen, Rufin VanRullen, Carl Michael Gaspar, Ye Zhang

## Abstract

Some neural responses are classified by the stimulus conditions leading up to that response while other neural responses are also classified by the morphology of the responses themselves. However, morphology-based classification may not be appropriate if one can nudge a neural response into looking like another neural response. Morphology-based classification occurs with the N170 and RP (Recognition Potential), ERP components that are studied in separate literatures and yet share much in common in terms of functionality. In this study, we demonstrate a gradual transformation in the morphology of the N170 to the RP using a simple parametric manipulation of forward masks that is unlikely to cause a change in the underlying processing. Both the N170 and RP are N1 components, meaning that they are the first negative deflection of the evoked response. However, the RP is often measured with a forward mask that ends at stimulus onset whereas the N170 is often measured with no masking at all. This study investigates how ISI may delay and distort the N170 into an RP by manipulating the temporal gap (ISI) between forward mask and target. The results revealed reverse relationships between the ISI on the one hand, and the N170 latency, single-trial N1 jitter (an approximation of N1 width) and reaction time on the other hand. Importantly, we find that scalp topographies have a unique signature at the N1 peak across all conditions, from the longest gap (N170) to the shortest (RP). These findings prove that the mask-delayed N1 is still the same N170, even under conditions that are normally associated with a different component like the RP. In general, our results suggest that greater caution should be taken to interpret the time course of a measured effect when forward masks are employed.

## 1 Introduction

Event related potential (ERP) components are important for inferring the time course of perceptual and cognitive processing (Carreiras, Armstrong, Perea, & Frost, 2014; Dien, 2009; Grainger & Holcomb, 2009). The most typical type of component, the kind studied in this paper, is a peak in the evoked response, which can be either a positive or negative deflection. Inference about processing can be relatively straightforward when components are classified almost exclusively by the stimulus conditions leading up to that component. In this case, delays and distortions in the component are interpreted as delays and distortions in processing. This is certainly the case with components like the readiness potential (Libet, Libet, Gleason, Wright, & Pearl, 1993) and the mismatch negativity (MMN) (Näätänen, Gaillard, & Mäntysalo, 1978). The mismatch negativity (MMN) is a response evoked by an unexpected stimulus in a sequence of regularly occurring stimuli. Importantly, an early MMN evoked by auditory stimuli is just as much an MMN as a late MMN evoked by visual stimuli (Winkler, Czigler, Sussman, Horváth, & Balázs, 2005)

However, confusion can arise when components are also classified by their morphology – their latency and shape. The importance of latency in defining some components is often reflected in component nomenclature like N170 (Shlomo Bentin, Allison, Puce, Perez, & McCarthy, 1996), which refers to a negative deflection occurring around 170 ms after an object from a familiar category appears. In some cases, peaks that are offset in time (i.e. with longer latencies in most cases) and slightly different in shape are thought to reflect different processes. While that interpretation is consistent with a cascade across processing levels, an equally valid interpretation of peak offset is a delay in processing. How can we decide between these two interpretations? This paper reports unique support for the delay interpretation in a comparison between 2 nominally different ERP components: the N170 and the recognition potential, or RP (Rudell, 1991; Martin-Loeches et al., 2001). The approach we take arises from the possibility that the presence of forward-masking during typical recordings of RP can lead to processing delay relative to the N170, which rarely involves forward masks. Masks are typically random patterns and so their mere presence is unlikely to involve a dramatic switch in processing. In order to test our delay-hypothesis we simply nudge the N170 into an evoked response that is morphologically equivalent to an RP, by using forward masks with progressively smaller ISI. Importantly, the experimental conditions at the extreme ends of our parametric manipulation match the conventional experimental designs of N170 and RP studies. The remainder of the Introduction describes N170 and RP in greater detail and motivates their comparison despite being studied in different literatures with very little cross talk. Experimental design is elaborated upon and more precise predictions are made in the context of the delay hypothesis.

The N170 is the first negative deflection in response to visually observed objects from a familiar category, like faces. It is described as being a critical temporal marker for the extraction of information from visually observed familiar objects (Martín-Loeches, 2007; Maurer, Zevin, & McCandliss, 2008; Tanaka & Curran, 2001). However, the latency of N170 can be influenced by various factors. For example, up-side down faces evoke an N170 that is delayed by about 10 ms, compared to that evoked by upright faces (Rossion et al., 1999). Another possible factor, which to our best knowlege has not been directly investigated, is inter-stimulus intervals (ISI). In a survey we made (Figure 1), one can see a monotonic relationship between N170 latency and ISI (Cao, Jiang, Li, & He, 2014; Dimigen, Kliegl, & Sommer, 2012; Fu, Feng, Guo, Luo, & Parasuraman, 2012; Harris & Nakayama, 2007; Kuehl, Brandt, Hahn, Dettling, & Neuhaus, 2013; Lin et al., 2011; Martens, Schweinberger, Kiefer, & Burton, 2006; Shen et al., 2017; Su, Mak, Cheung, & Law, 2012). In fact, a very short ISI can delay the N170 up to ∼80 ms later than the typical latency, which is an enormous effect compared to the face inversion effect. Moreover, if a study uses very short ISI between forward masks and targets (for example, Su et al., 2012; studies marked with orange color in Figure 1), we can see that not only is the latency delayed, but also the waveform is wider and deformed enough to make peak isolation challenging. Taken together, the results drawn from these separate studies suggest that the N170 can be nudged forward and widened by the recency of prior stimulation. The most extreme delay should occur with zero ISI (e.g., Su et al., 2012), at which point the N170’s morphology might be indistinguishable with another ERP component: the recognition potential (RP).

**Figure 1.**
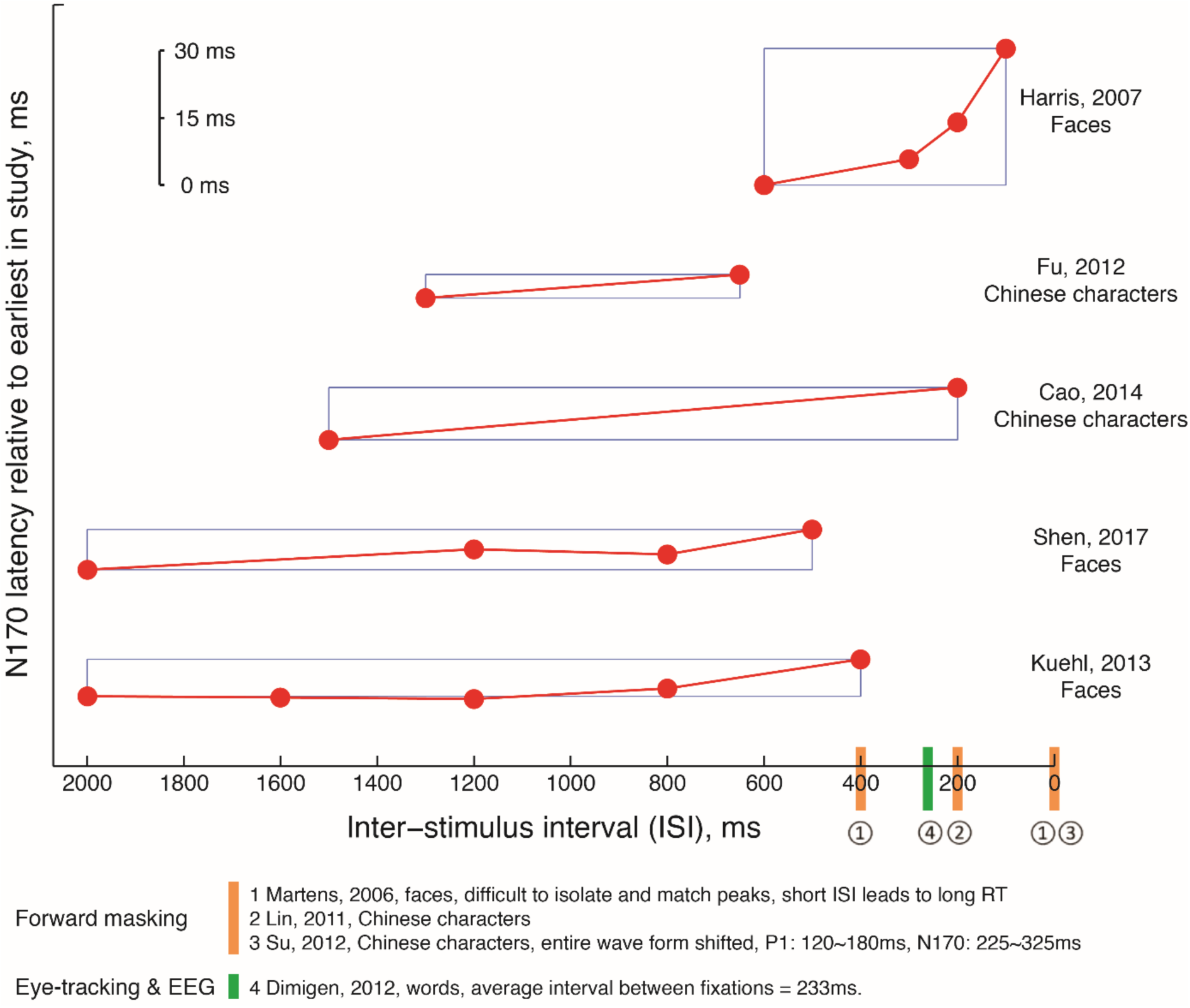
Summary of studies that used short ISI. The studies marked in yellow are forward-masking studies. These studies provide more limited results than the adaptation studies shown above the x axis, because: they either have only a single ISI, they do not report N170 latencies, or those latencies are difficult to estimate from the figures because the grand-average waveforms are complex. The study marked in green is an eye-tracking & EEG study, the average interval between reading fixations during a combined EEG and eye-tracking study on reading can be consider like an “ISI”.

RP is usually studied in a totally separate body of literature focusing on the time course of word recognition (Dien, 2009; Martín-Loeches, 2007). Unlike the N170, the RP is usually measured with a paradigm called rapid stream stimulation (RSS) which presents multiple masks before a target in succession and multiple masks after. However, RP is similar to N170 in many other aspects. Primarily, both the RP and N170 are responses to the visual presentation of familiar objects. And there are also a number of morphological similarities between these two ERPs. Both of them are almost always N1 components (for an non-N1 RP, see Rudell, 1991; 2010). Both ERPs are observed at similar channels and have similar topographies (S. Bentin, Mouchetant-Rostaing, Giard, Echallier, & Pernier, 1999; Hinojosa, Martín-Loeches, Gómez-Jarabo, & Rubia, 2000; Martín-Loeches, Hinojosa, Fernández-Frías, & Rubia, 2001; Martin-Loeches, Hinojosa, Gomez-Jarabo, & Rubia, 2001; Maurer, 2008; Maurer, Brandeis, & McCandliss, 2005; Maurer et al., 2008). The one major morphological difference between N170 and RP is delay: the RP typically occurs around 250 ms post-stimulus, which is longer than the typical latency of N170. And because of the delay, RP is often linked to higher level processing stages than the processing stage revealed by N170 (Dien, 2009). However, this widely-accepted interpretation of RP does not consider the presence of forward-masking during typical recordings of RP. If the forward mask merely delays the brain signal in response to the target, could the RP just be a delayed brain response such that its mask-free latency would actually be ∼170 ms? To put it simply, is RP actually an N170 delayed by forward masks?

Taken together, our study aims to answer two questions: 1) Is the N1 systematically delayed and warped by ISI? 2) Is RP a delayed version of N170? Answers to these questions can only be obtained by establishing the relationship between ISI and N170. In this study, we measure electrophysiological responses evoked by rapidly displayed Chinese characters, and we focus mostly on characteristics of the N1 and vertex positive potential (VPP). VPP is measured at a vertex channel such as Cz and is considered to reflect the same process the N170 does (Joyce & Rossion, 2005). Our subjects also performed a 1-back task on Chinese characters, and we measured their reaction times. In these respects, our study is like many others in both the N170 and RP literatures. However, our study is unique in three ways: First, across 14 conditions, we parametrically manipulate the temporal proximity between the target and a forward-mask image, from 0-gap (quasi-RP condition; gap denotes ISI) to infinity-gap (or no-mask, the typical N170 condition). Second, we focus on the specification of waveform-morphology in order to determine whether the N1 component obtained in N170 and gap conditions is the same. Finally, we obtain lots of data for each subject (150 trials for each of the 14 conditions). Altogether, we are able to obtain unique “gap-metric” functions, the ERP-component equivalent of a psychometric function for 3 major aspects of N1 morphology: N1 latency, N1 single-trial jitter, and topographical similarity to the N170. In addition, reaction times measured across gap conditions also allows us to assess the functional consequences of gap-related waveform changes. Our experimental design allows us to isolate the effect of a single masking stimulus on variability in N1 morphology. All these allow us to answer the first question and part of the second question. Considering that applying only one forward mask per trial is not typical in the majority of RP studies (but see: Dien, Brian, Molfese & Gold, 2013), to completely answer the second question, typical RP was also measured with the RSS paradigm as a control condition and compared with all the other 14 conditions.

## 2 Materials and methods

### 2.1 Participants

Nine healthy adult participants (all right-handed, age ranging from 24 to 27, 4 males) with normal or corrected-to-normal vision took part in the study and were paid for their participation. Data from two participants were rejected due to too many artifact-contaminated trials (e.g., eye blinks, head movements and α wave). Valid data from seven participants were reported here. All participants were native Chinese who had more than 10 years of education in speaking and reading Chinese characters. Written informed consent was obtained from each participant prior to the experiment.

### 2.2 Stimuli

Stimuli were gray scale images of 150 Chinese noun characters and 150 masks. Chinese characters were chosen from the Modern Chinese Corpus of Center for Chinese Linguistics (Peking University, 2015) (see http://ccl.pku.edu.cn:8080/ccl_corpus/index.jsp?dir=xiandai). The frequency of each character was higher than 105 occurrences per million in this corpus, mean frequency of all characters is 250.95 occurrence per million. The number of strokes in Characters ranges from 7 to 12 (mean 9.18). The masks were made up of the 150 Chinese characters by cutting these characters into ‘16’ portions vertically and randomly permuting their horizontal position. These masks and Chinese characters were matched in visual attributes. The size of the basic set of characters and masks were around 4.4°×4.4° of visual angle.

### 2.3 Procedure

After electrode-cap placement, participants were seated in a dim light and sound-attenuated shielded room, at a viewing distance of 60 cm from the computer monitor. Stimuli were presented with MATLAB (MathWorks, Natick, Mass) using the Psychophysics Toolbox (Version 3.0.12; http://psychtoolbox.org) (Brainard, 1997; Kleiner et al., 2007) on a light gray background.

The sequence of events for each trial, summarised in Figure 2, is as follows: In each trial, a “+” fixation was presented at first in the center of the screen for 500 ms, which was followed by a blank for 800 ms. Then, a forward mask, a character, and a backward mask were presented orderly, each for 250 ms. Each trial was ended by a blank of 800 ms. The masks in each trial were randomly chosen from all the masks. Participants were instructed to make a response by pressing the ‘m’ key as quickly and accurately as possible when a repeated character was detected in successive trials. The ISI (i.e., temporal gap) between the preceding mask and the character varied from 0 to 600 ms with a step of 50 ms. So totally, there were 13 gap conditions. Additionally, there were two more conditions, one was without the preceding mask in order to elicit the classic N170 (no-mask condition), in which the blank interval between the fixation and the character was 1050 ms; for the other one the RSS paradigm was used, in which 3∼5 masks were presented prior to the targets and 4∼6 masks were presented after to elicit typical RP (RSS condition), which served as a control condition. The stimulus presentations for different conditions are described in Figure 2. All 15 conditions were separated by blocks; each block had only one condition. There were 58 trials in each block where 8 characters were randomly chosen to be repeated ‘targets’ (same character as shown on previous trial). Each condition was tested with three blocks, totally 174 trials for one condition. The experiment started with a practice block of 58 trials (these stimuli were not used in the main test). The order of blocks and trials in each block were randomised, and the RSS condition (control) was conducted after the subjects completed all the other conditions.

**Figure 2.**
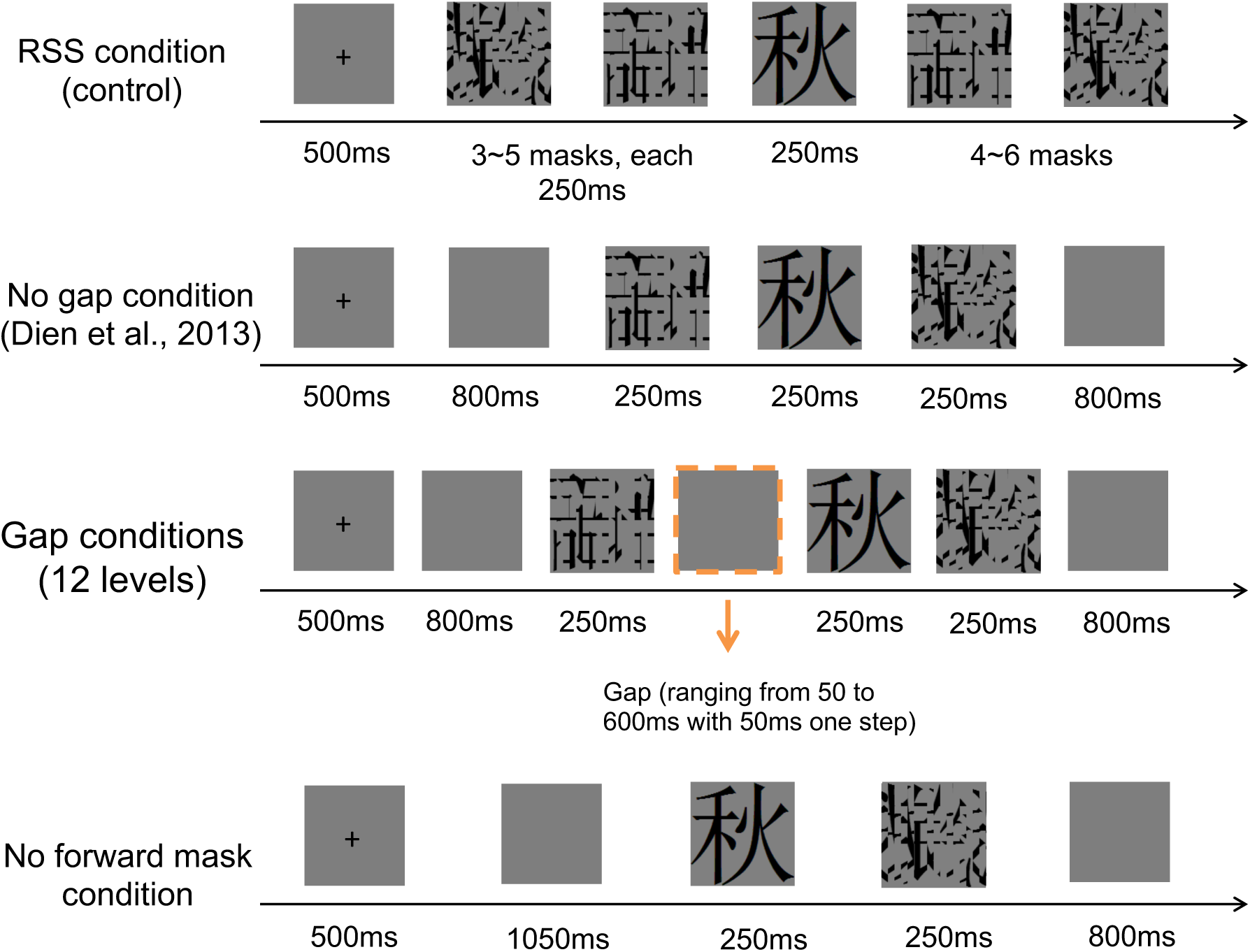
Trial sequence for each condition. The sequence shown at the top is for the RSS (typical RP) condition, the second sequence is for the 0-gap (quasi-RP) condition. The sequence shown at the bottom is for the no-mask (N170) condition. And the sequence in the middle shows how intermediate gap conditions are generated by varying the duration of the central event (a blank screen, outlined with a dashed line in this figure). Note how the overall timing is equated between the 0-gap and no-mask conditions.

### 2.4 EEG recording and preprocessing

Brain electrical activity was recorded from 32 scalp sites using tin electrodes mounted in an elastic cap (BrainAmp, Brain Product). Vertical and horizontal eye movements were monitored using two electrodes placed on the outer canthus of right eye and in the inferior areas of the left orbit. The ground electrode was placed along the midline, ahead of Fz, and impedances were systematically kept below 7 kΩ. Signals were digitised at a sampling rate of 500 Hz and band-pass filtered at 0.016∼70 Hz. Potentials were referenced on-line to the FCz electrode and averaged-referenced off-line. Participants were asked to minimise eye movements, head movement, and swallowing during the recording.

EEG data was low-pass filtered at 30 Hz and high-pass filtered at 0.1 Hz. Data was epoched from 1000 ms before the onset of stimulus to 1000 ms after the onset of stimulus. Only non-repeated trials were extracted. Artefact rejections were applied over the epoched data. Trials with α wave and abnormal trend and blink during the presentation of stimuli were rejected by visual inspection. After trial rejection, the remaining numbers of trials did not vary much across conditions (see Table 1).

**Table 1.** Number of trials per condition, summarised by quartiles across subjects. Q1 to Q3 refer to the first to third quartiles; i.e., Q2 is the median.

|  | RSS | 0 | 50 | 100 | 150 | 200 | 250 | 300 | 350 | 400 | 450 | 500 | 550 | 600 | N170 |
| --- | --- | --- | --- | --- | --- | --- | --- | --- | --- | --- | --- | --- | --- | --- | --- |
| Q1 | 126 | 125 | 124 | 128 | 128 | 125 | 133 | 121 | 119 | 122 | 118 | 107 | 124 | 126 | 114 |
| Q2 | 138 | 136 | 136 | 136 | 134 | 136 | 138 | 134 | 138 | 139 | 134 | 142 | 140 | 140 | 133 |
| Q3 | 141 | 139 | 142 | 145 | 138 | 142 | 145 | 143 | 143 | 144 | 137 | 144 | 145 | 143 | 143 |

### 2.5 Analysis

#### 2.5.1 N1 latencies (mean ERP)

The peak latencies and amplitudes of N1 were automatically extracted at the maximum (negative) amplitude value between 130 and 330 ms at P7 electrode (left occipio-temporal area), and the peak latencies and amplitudes of VPP were automatically extracted at the maximum (positive) amplitude at Fz electrode (fronto-central area) within the same time window. Automatic extraction was performed using ERPLAB (Lopez-Calderon & Luck, 2014) and EEGLAB functions (Delorme & Makeig, 2004).

#### 2.5.2 N1 widths (single-trial jitter)

N1 component width, measured directly on the mean ERP waveform, is difficult to define. Some subjects lack distinctive peaks around the N1 (P1 and P2) to serve as delimiters. Also, subjects vary in the shape of their N1 component; some having more kurtosis than others, and others being more asymmetrical. Therefore, we measure single-trial N1 components and take the standard deviation of their latencies across trials - their temporal jitter - to be a proxy measure of component width. Several studies have used this method to establish reliable differences in N1 latency variability across experimental conditions (e.g., Navajas, Ahmadi, & Quian Quiroga, 2013; Zhang, Luo, & Luo, 2013, and across subject populations (Milne, 2011). Single-trial N1 peaks were estimated from 150 ms windows centred on the latency of the N1 in the mean ERP of the appropriate gap condition and subject. Within each window of each trial, we used custom Matlab scripts and publicly available Matlab code to estimate the latency of the most negative local peak (findpeaks.m, authored by Professor Tom O’Haver at University of Maryland, Version 6.0, Last revised March, 2016; http://terpconnect.umd.edu/~toh/spectrum/findpeaks.m).

#### 2.5.3 Decay function relating N1 latency and jitter, to gap

We used the following exponential function to relate both N1 latency, and N1 jitter, to gap, separately for each subject:

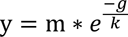

where y can be either N1 latency (ms), or N1 jitter (standard deviation, ms), g is gap (ms), m specifies the starting point (y at gap 0), and k is the inverse of the decay rate (small k means fast decay). Best fitting parameters were obtained by minimising the sum-of-squares error using Matlab’s fminsearch.m (Matlab R2014a).

#### 2.5.4 Statistical analyses of correlations

For statistical analyses of correlations between N1 latency and RT, and between N1 latency and VPP latency, we used the lmer program of the lme4 package (Bates, Mächler, Bolker, & Walker, 2014) for estimating fixed and random coefficients (subjects). Our focus is on the coefficient between N1 latency and RT, or between N1 and VPP latencies, modelled as fixed effects in separate models. Variation in intercept across subjects is modelled as a random effect; hence the use of a linear mixed model. Our main results concern the coefficient associated with our fixed effects, which is evaluated in two ways. First, we examine an anova-based model comparison between a subjects-only model and the subjects-plus-coefficient model (the full model). Second, and more importantly, we use a parametric bootstrap of the full model to generate a confidence interval for the coefficient. This package is supplied in the R system for statistical computing (The R Development Core Team, 2011). For a comparison with more traditional methods, we also include the results for repeated-measures multiple regression analysis (rmMRA). Performing an rmMRA is simple: Multiple linear regressions are performed separately for each subject, then a t-test can be performed to determine if the mean of a coefficient value across subjects, is significantly different from zero.

## 3 Results

### 3.1 Gap-metric functions

Our main data are mean ERP for each condition and each subject. A nice visual summary for each subject can be constructed by what we call a ‘stack plot’: ERP stacked vertically across conditions (in order of gap), and colour-coded for amplitude. The stack plot for subjects 1 is shown in Figure 3 (code for producing this stack plot is avalaible on Github: https://github.com/SourCherries/firecracker. Stack plots for the remainder of subjects can be found in supplementary Figure 1). A large time-course, from −800 to 600 ms is shown, so that one can see responses to both masking and target stimuli. The main observation is a systematic change in shape and overall timing from the N170 condition to the widest gap, and then to smaller and smaller gaps: as a mask is introduced and its temporal proximity to the target increases, the neural response to the target appears to delay and widen proportionally. We can quantify these relations more precisely by examining the relation between gap (ms) and N1 peak latency (ms), N1 single-trial jitter (SD ms), and reaction time (ms).

**Figure 3.**
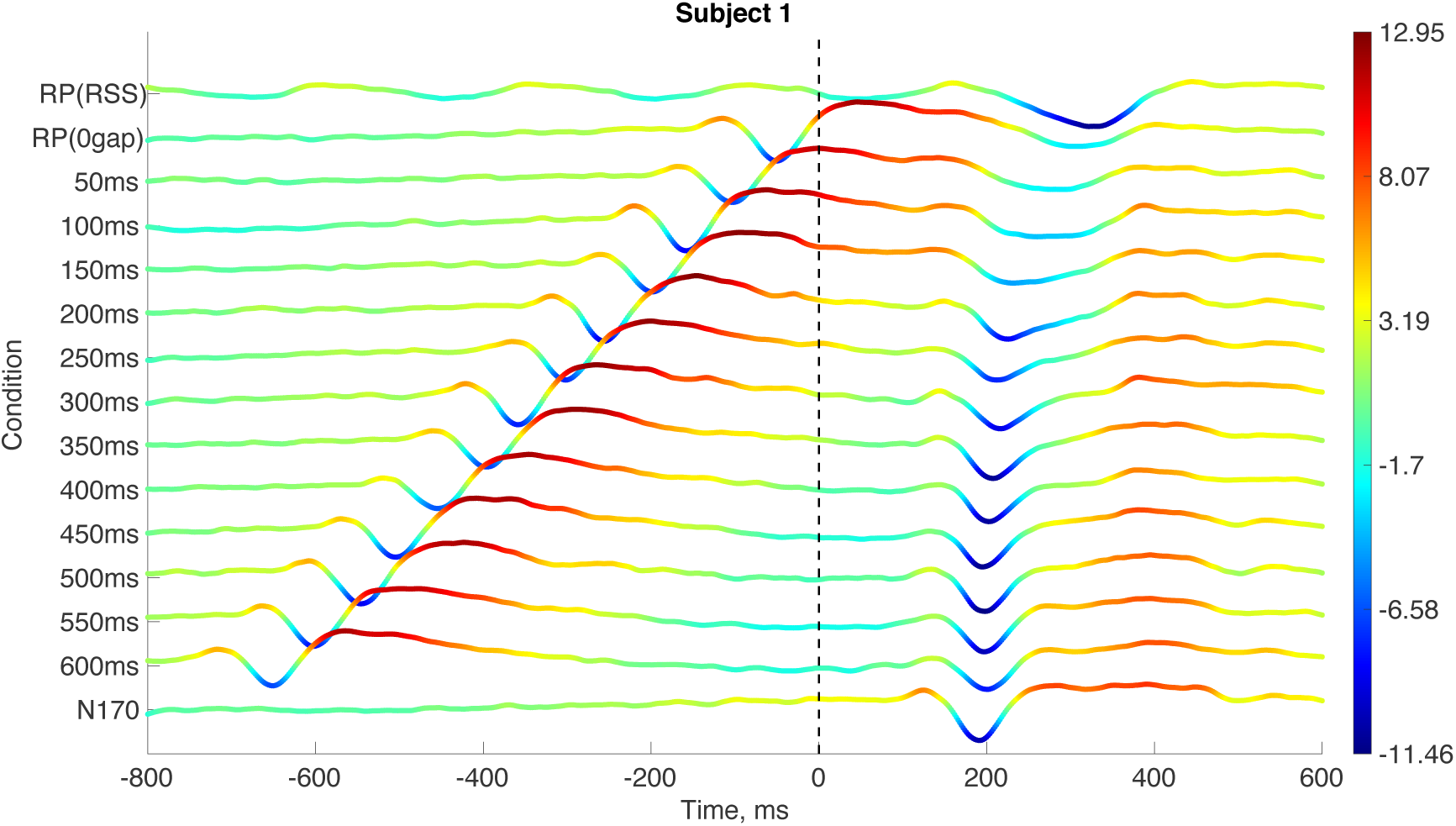
ERP stack plot for subject 1 (top). Mean ERPs are stacked vertically across conditions and voltage is colour-coded, with a corresponding colour-bar to the right.

Each gap function (N1 latency, jitter and RT), measured separately for each subject, is normalized by its respective value for the N170 condition. We do this because the N170 does not have a ‘gap’, since there is no mask in that condition (arguably, ‘gap’ is effectively infinity). Normalization is also a way to easily check if values are different from their natural baseline (N170): latencies should be mostly above 0, as with jitter values and RT. For N1 latency and RT, N170 latencies are subtracted. And for jitter, which are in units of standard deviation (ms), we take the log of jitter divided by N170-jitter.

Gap-metric functions for N1 latency, jitter, and reaction time, are shown in Figure 4. For each subject, N1 latencies and jitters are a clear exponential function of gap. There are also clear individual differences in decay rate, so we order gap functions for latency and jitter from subjects with the smallest to the highest decay rates (from high k to small k), which are related by a power function almost perfectly between latency and jitter (Figure 5). Therefore, N1 jitter and latency decay ‘together’.

**Figure 4.**
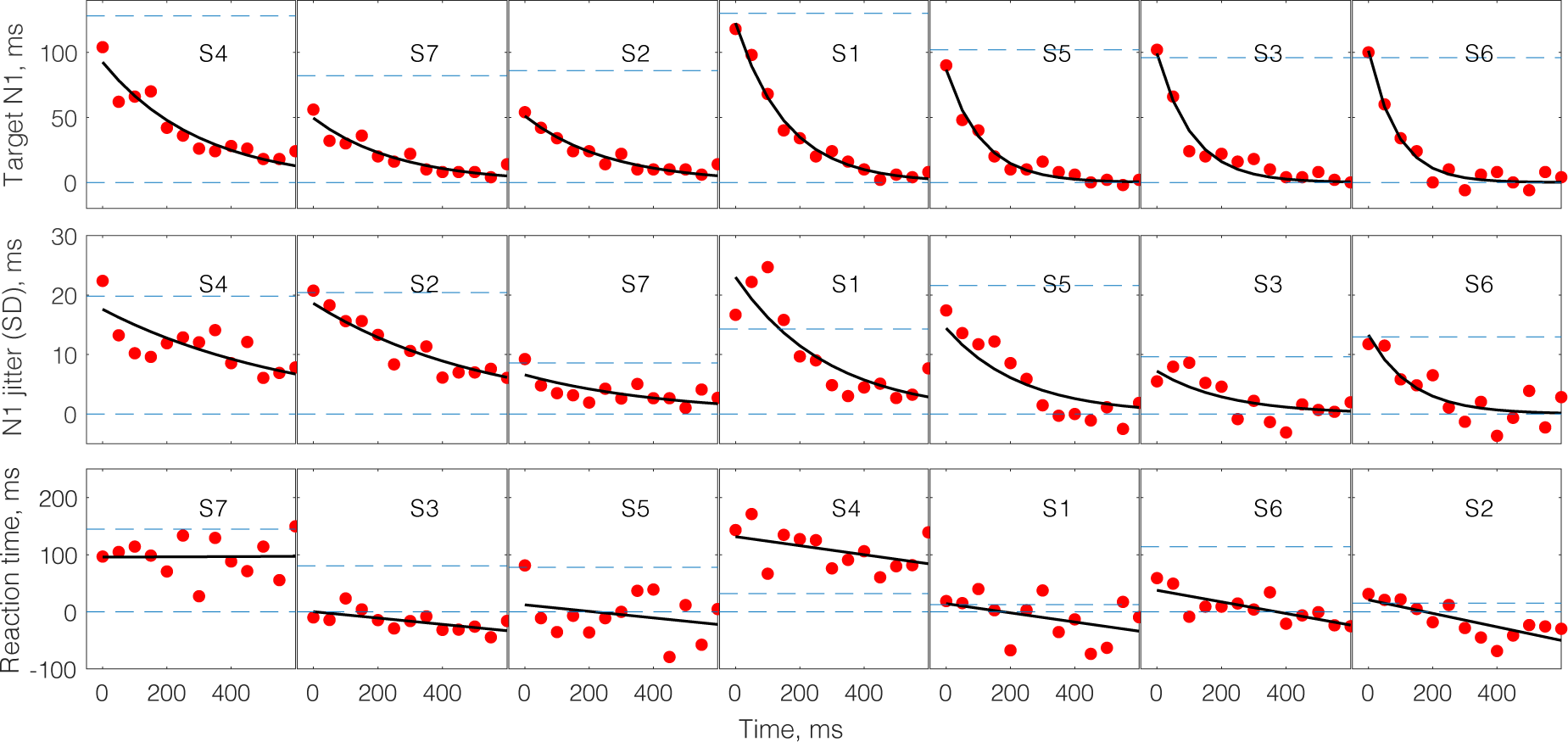
Gap functions for N1 latency, N1 jitter and reaction time, from top to bottom. The upper dash lines represent RP (RSS condition), and the bottom dash lines represent N170 (no forward mask condition).

**Figure 5.**
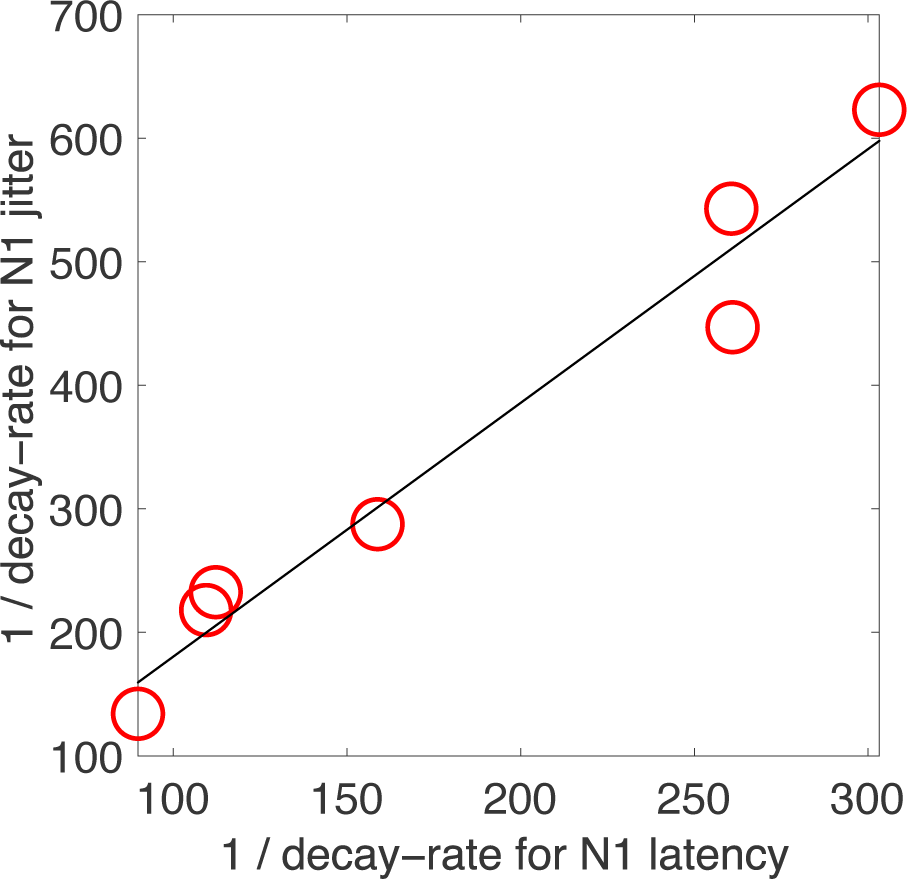
Decay rates of N1 latency and jitter are related by a power function. The fitted line is k_n1_jitter = m*k_n1_latency^gamma, where best fit parameters are m=0.0884 and k = 1.6315.

Figure 4 also shows a clear trend for an inverse relation between RT and gap (lines are fit here instead of exponential functions, and subjects are ordered by intercept). RT is more closely examined in the next section.

To further confirm that the jitters are the main factor that widen the target N1s, we removed the variance of the single trial latencies by aligning the single-trial peaks then averaged the trials that belong to each condition. All the average N1s appeared to have similar width after removing the single trial jitters (Supplement Figure 2).

In agreement with the studies in our survey (Figure 1), our results (Figure 3 and 4) show that the forward-mask changes the timing of the N1 waveform, depending on how close it is to the target image: the smaller the gap, the greater the change in N1 (both delay and widening). If that is true, then one would also expect that the speed of a subject’s own neural response to the mask - as an intermediate event between mask and target - would have a similar effect: if their mask-N1 is very slow (long latency) then their ‘effective gap’ is much smaller than the ‘nominal gap’ value; but if their mask-N1 is very rapid (short latency) then their ‘effective gap’ is closer to the ‘nominal gap’ between mask- and target-image onsets. Therefore, individual differences in mask-N1 latency should accurately predict individual differences in target-N1 decay rates. As expected, individual differences in decay rate, for both N1 latency and jitter, are excellently predicted by the latency of a subject’s N1 response to the masking stimuli (Correlation between Mask N1 latency and parameter k of target N1 latency: r = −0.8583, p = 0.0134; Correlation between Mask N1 latency and parameter k of target N1 jitter: r = −0.9118, p = 0.0042) (Figure 6).

**Figure 6.**
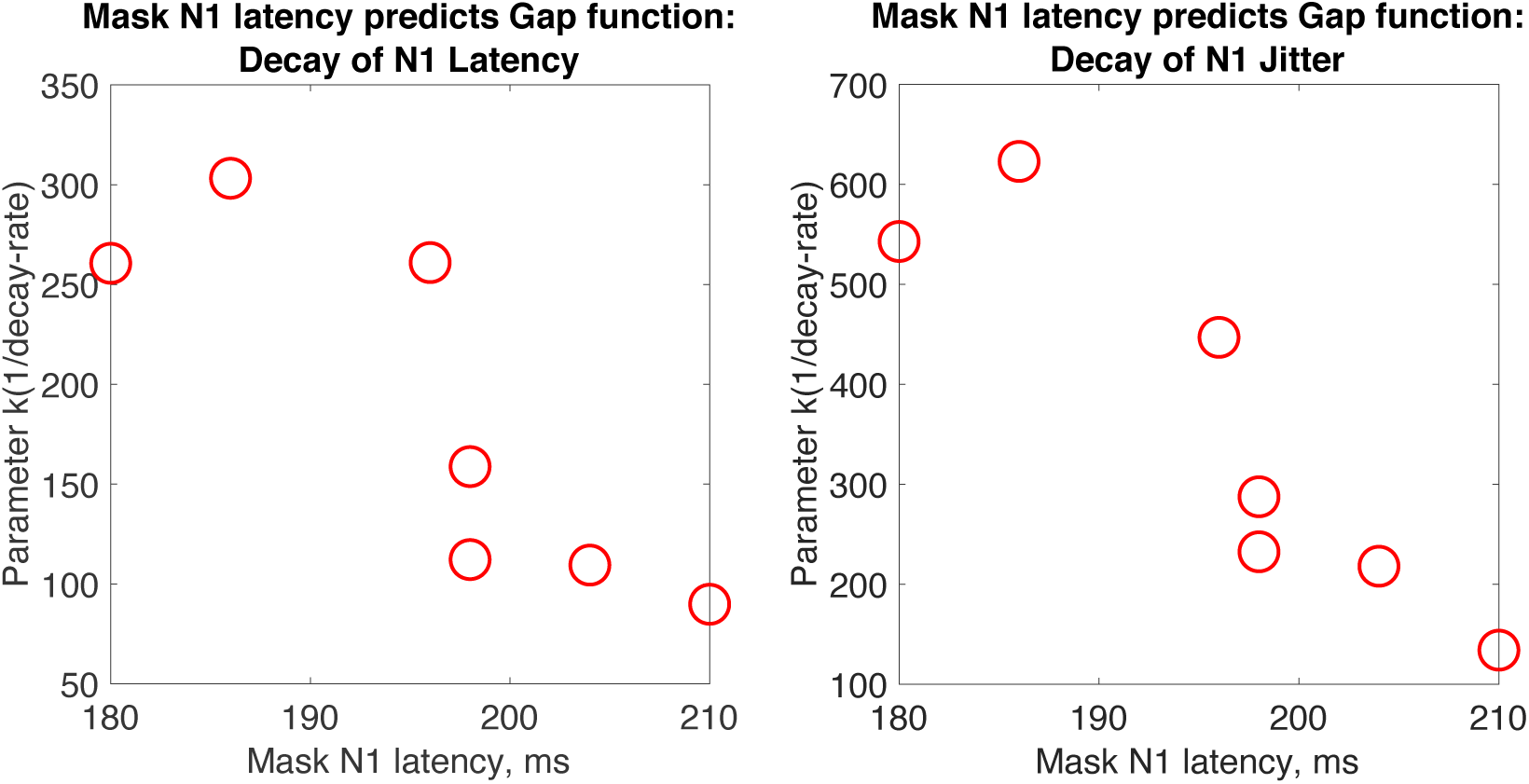
Latency of N1 response to masking stimulus predicts k-parameter (1/decay-rate) of target-N1, for both N1 latency and N1 jitter.

### 3.2 N1 latency and RT

We employ both a linear mixed model (LMM) and repeated-measures multiple-regression analysis (rmMRA). For LMM, we perform 2 model comparisons. First, we compare subject-intercept only, versus subject-intercept plus N1-latency, with reaction time (RT) as the dependent variable. These models are not nested; they have different fixed-effect structures. Therefore, we estimate these models using maximum likelihood estimation rather than restricted maximum likelihood (Knoblauch & Maloney, 2012). A LMM without N1-latency fits significantly worse than the complete model, with a delta-Chi-square (1 d.f.) = 20.55, p = 5.798e-06, for the decrease in log likelihood. Similarly, rmMRA results (summarised in Table 2, along with result for the full LMM model), demonstrate a significant N1-latency and RT correlation. An additional nested, model comparison, that examined the effect of including random slopes was insignificant, with a delta-Chi-square (1 d.f.) = 0.96, p = 0.33. Evaluation of the N1-latency and RT correlation can be made on the significance of the LMM model comparison, and the rmMRA. However, p-values obtained from model comparisons of LMM (Knoblauch and Maloney, 2012), and from rmMRA (Baayen, Davidson, & Bates, 2008), tend to be liberal, so we focus mainly on the confidence intervals of effects, which are based on a parametric bootstrap of 1000 samples from our model based on N1-latency and random subject intercepts (using restricted maximum likelihood). The 95-percent confidence interval for the coefficient for N1-latency is [0.344, 0.845]. The point estimate (shown in Table 2) is 0.59, so for every 10 ms delay in N1, we expect a delay in reaction time of about 6 ms.

**Table 2.** Summary of fixed effects for both LMM (using restricted maximum likelihood) and rmMRA models of RT, predicted by N1-latency. Additionally, for the LMM, we have a random effect (not shown in this table): the standard deviation associated with subject (intercept) is 61.44, which ends up being approximately the same as the SD of intercepts measured with rmMRA. t>2.4 is significant at 5% level (two-tailed).

|  | LMM |  |  | rmMRA |  |  |
| --- | --- | --- | --- | --- | --- | --- |
|  | Coefficient | SE | t | Coefficient | SE | t |
| Mean RT, ms | 524.84 | 35.32 | 14.86 | 503.06 | 23.65 | 21.27 |
| N1 latency, ms | 0.59 | 0.12 | 4.81 | 0.71 | 0.14 | 4.95 |

### 3.3 N1 and VPP latencies

In the N170 literature, the VPP component is sometimes measured alongside the N170 component. Joyce and Rossion (2005) find that the amplitudes of the N170 and VPP components vary in a precisely inverse manner and the peaks of the N170 and VPP are temporally coincident. If the N170 latencies were delayed by the forward mask, VPP latencies should also be delayed along with the N170 latencies; as with the timing of every pair of N1 and VPP in each gap condition. In other words, the N170 latencies should correlate with the VPP latencies, as shown by the LMM below.

For LMM, first we compare subject-intercept only, versus subject-intercept plus VPP-latency. We estimate these models using maximum likelihood estimation. A LMM without VPP-latency fits significantly worse than the complete model, with a delta-Chi-square (1 d.f.) = 173.7, p = 2.2e-16, for the decrease in log likelihood. Similarly, rmMRA results (summarised in Table 3, along with result for the full LMM model), demonstrate a significant VPP-latency and N1-latency correlation. An additional nested, model comparison, that examined the effect of including random slopes was insignificant, with a delta-Chi-square (1 d.f.) = 2.06, p = 0.15. Evaluation of the VPP- and N1-latency correlation can be made on the significance of the LMM model comparison, and the rmMRA. Focussing on our LMM of N1-latency, based on VPP latency and random subject intercepts, we perform a parametric bootstrap of 1000 samples (using restricted maximum likelihood). The 95-percent confidence interval for N1-latency is [0.79, 0.94].

**Table 3.** Summary of fixed effects for both LMM (using restricted maximum likelihood) and rmMRA models of N1-latency, predicted by VPP-latency. Additionally for the LMM, we have a random effect (not shown in this table): the standard deviation associated with subject (intercept) is 9.2, which ends up being slightly higher than the SD of intercepts measured with rmMRA (8.8). t>2.4 is significant at 5% level (two-tailed).

|  | LMM |  |  | rmMRA |  |  |
| --- | --- | --- | --- | --- | --- | --- |
|  | Coefficient | SE | t | Coefficient | SE | t |
| Mean N1 latency, ms | 31.521 | 8.944 | 3.524 | 34.78 | 17.081 | 2.04 |
| VPP latency, ms | 0.862 | 0.038 | 22.458 | 0.84 | 0.082 | 10.32 |

### 3.4 Topographies

Analysis described here measures matches in topographical distributions between time frames, and between conditions. Inner-products were used instead of (normalised) correlations because measuring correlation would require centring and normalising each topography on a frame-by-frame basis. However, such normalisation, we argue, would throw away information about changes in polarity across time, as well as overall voltage energy.

Assuming the N1 topography from the N170 condition is a unique marker associated with processing at N1 (Maurer et al., 2005), the strength of its inner-product with topographies from other gap conditions should roughly predict the timing of the N1 in those conditions. How well are N1 latencies in each gap condition predicted by their topographical match with the N170 condition at N1?

In each gap condition, we measured the time course of inner-products between the N1 topography from the N170 condition to all other topographies in that condition. If the N1 peak is similar across gap conditions, in the sense that it reflects the same process that underlies the N1 in the N170 condition, then latency of the maximum inner-product along the time-course for a given gap condition should roughly match the timing of the N1 in that condition. In addition, this logic also applies to the RSS condition. To be clear about this analysis, we include markers in Figure 7 that depict both the timing of N1 latency (black triangles), and the timing of the max inner-product with N170 topography (red, upside down triangles). One can clearly see that maximum latencies follow N1 latencies across gap condition. To precisely quantify this correlation for every subject, we include a scatter plot between maximum-inner-product and N1 latencies in Figure 8, using different coloured markers for each of the 7 subjects. This plot is convincing: across gap and subject, topographical distributions across time are most similar to the N170 topography at the latency of the N1 (r = 0.9424, p = 8.7283e-51).

**Figure 7.**
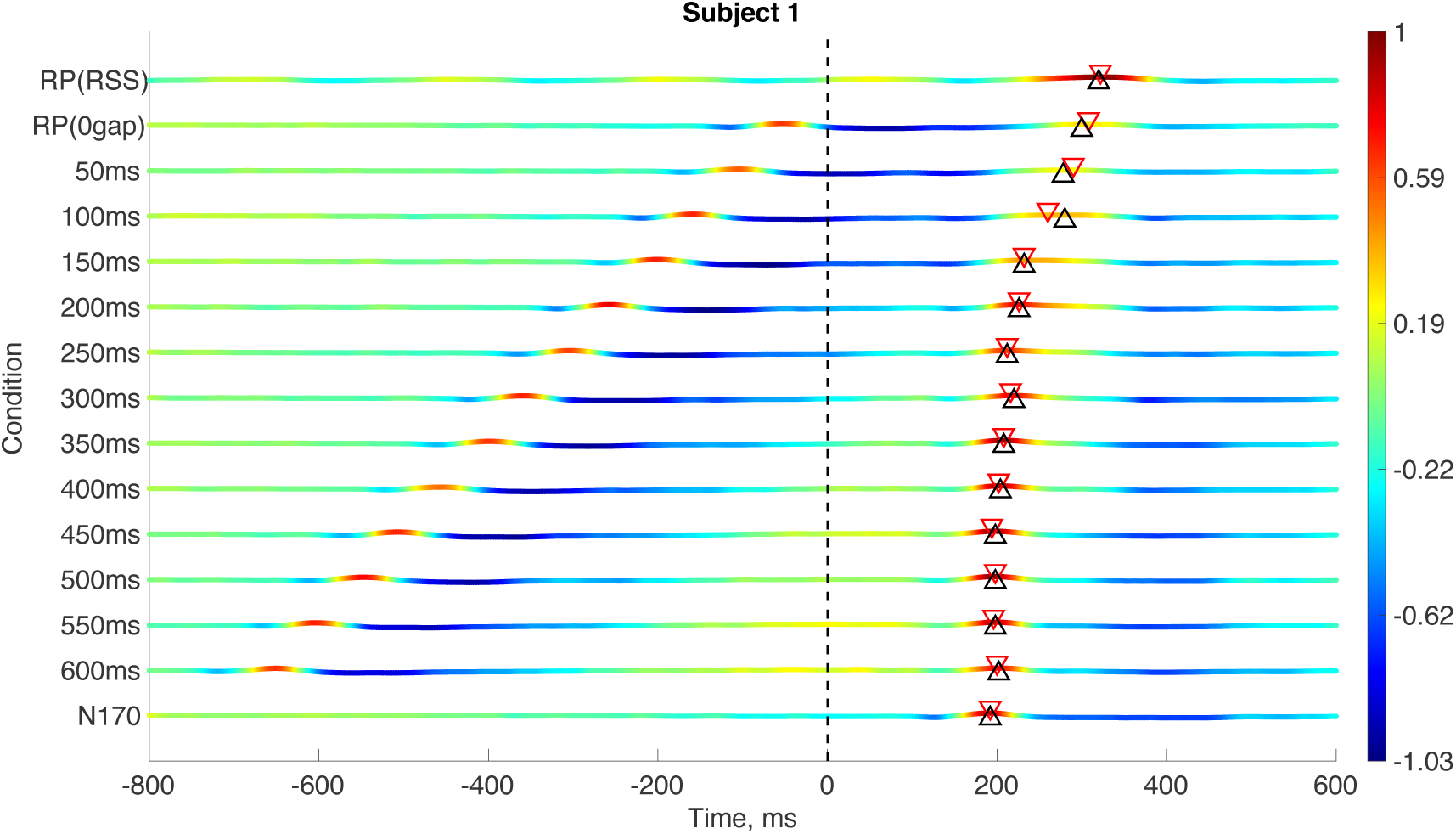
Time-course of inner product between N170 topographical distribution (at N1 latency), and topographies at every other time-frame, and condition. To improve visualisation, each inner-product time-course is normalised by its maximum inner-product. In each condition, a black triangle (pointing up) shows the latency of the N1 for the mean ERP in that condition; and a red triangle (pointing down) shows the latency of the topography in that condition which has the highest inner-product with the N170 topography (at N1), within 0 to 400 ms. According to our hypothesis, black and red triangles should roughly match up.

**Figure 8.**
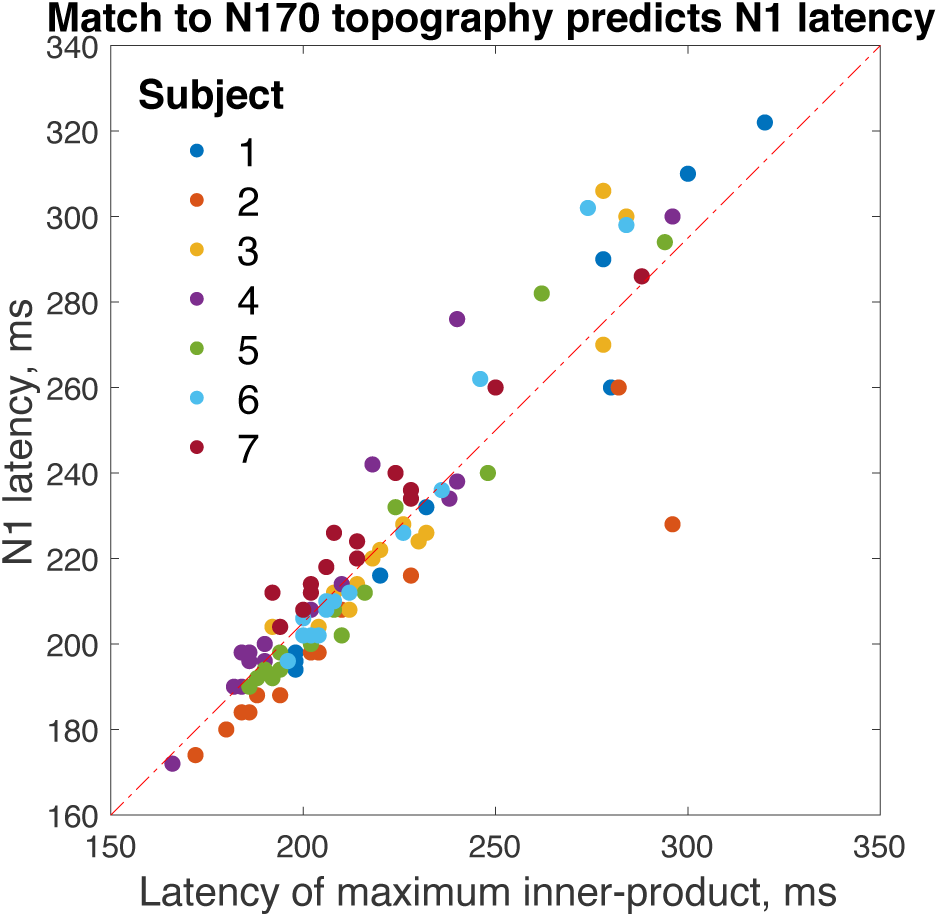
The latency of maximum inner-product with N170 topography (at N1) for each gap condition, predicts the N1 latency for that condition, for each subject. For clarity, note that, the x-axis here corresponds to the black triangles depicted in Figure 7 (for subjects S1); while the y-axis corresponds to the red triangles in that Figure.

A same analysis was done to match the RP (RSS condition) topography at RP latency with every frame of every condition. The question is similar: are topographies in each gap condition most similar to the RP topography at their respective N1 latencies? The answer to the question is yes. The results using RP topography are basically identical with the results using N170 topography; the RP version of figure 7 and 8 can be found in the suplementary material (Supplementary Figure 4 and 5)

## 4 Discussion

The current study tested the hypothesis that the N170 can be nudged forward by forward-masking into a component that is morphologically equivalent to an RP. All aspects of waveform morphology revealed a full picture of how ISIs and forward masks influence the target N1 and therefore provided compelling evidence that: (1) The delayed N1 can be recognized as N170; and (2) RP is a delayed version of N170. For each subject, N1 latency fell from its highest level in the 0-gap (quasi-RP) condition, toward the asymptotic level given by the latency in the no-mask (N170) condition. Individual differences in the decay parameter were well predicted by how quickly their mask-evoked N1 occurred: Faster N1 responses to the mask resulted in slower decay rates. Single-trial N1 jitter, our approximation of N1 width, behaved in the same manner as N1 latency: for each subject, an exponential decay with gap increase; and decay parameters well predicted by latency of mask-evoked N1. Target N1 of all the conditions had similar width after removing single-trial N1 jitter. Importantly, reaction times were also delayed by gap reductions, and were significantly correlated with N1 latency, consistent with the idea that a delayed N1 signals a delay in information processing. Finally, we demonstrate that the specific topographical distribution of voltage at N1, measured in the N170 condition, is both unique to the N1 time-point in that condition, and best matched to a small temporal window around the N1 time-point, for every gap condition and RSS condition (typical RP): 1) topographical similarity to the N170 (at N1) follows the delay in N1 latency as the gap between mask and target narrows. 2) topographical similarity to the N170 in the RSS condition appears at same time as the latency of the typical RP.

### The delayed N170 should be categorized as N170

The finely-spaced gap conditions spanning 0∼600 ms allowed us to uncover a continuum of changes in N1 latency and peak width. In agreement with the results of the studies using the short ISIs sumarised in our introduction, our results showed that target N1 was delayed when a short ISI was used. Moreover, the width of target N1 became wider and noisier when the gap is short, in line with Su et al. (2012). Both latency delay and the wider width can be explained by short ISIs and forward masks using an exponential decay function, and the decay rate can be predicted by the mask N1 latency. When we removed the jitter and aligned the single-trial peak, the N1 widths of different conditions were actually similar (Supplementary Figure 2), indicating that the single-trial jitter was the main cause of the widened width of N1.

In addition to the studies summarized in Figure 1, of which 4 sudies utilized adaptation (Cao et al., 2014; Fu et al., 2012; Kuehl et al., 2013; Shen et al., 2017), many other N170 studies have adopted short ISI, such as studies using priming (Campanella et al., 2000; Jemel, Pisani, Calabria, Crommelinck, & Bruyer, 2003) and rapid serial visual presentation (RSVP)(Gao et al., 2011; W. Luo, Feng, He, Wang, & Luo, 2010; W. Luo et al., 2013; Robinson, Plaut, & Behrmann, 2017). If the N1 latency was out of the typical N170 time window (as almost all these RSVP studies have shown), these studies would have faced one problem, that is: when did the measured effect actually happen? At ∼170ms where the typical N170 occurs? Or at the actual latency the effect was measured? This question is difficult to answer based only on the results of these prior studies. Even if the topography of the N1 seemed similar with the topography of a typical N170 (Gao et al., 2011; W. Luo et al., 2010, 2013), there was no evidence supporting the idea that the N1 was merely pushed forward by the previous stimulus, and that the topography was not by chance akin to N170. Now our current study has provided this missing evidence.

Given that both the signal of the whole brain and the behavioral responses were delayed by the forward masks, we can conclude that the whole information process was delayed. One plausible expalanation for the delay could be that it was the result of the refractory period of the visual system (Berry & Meister, 1998; M. C. Teich, Matin, & Cantor, 1978; Malvin Carl Teich & Diament, 1980). Before processing the next stimulus, neurons require time to recover from processing the previous, which is compatible with the inverse correlation between mask N1 latency and the parameter k (1/decay rate) (Figure 6), suggesting that the slower the brain responds to the forward mask, the more the target N1 latency is affected. This explanation based on the refractory period could hold for not only the paradigms using forward masks, but also the paradigms presenting any kinds of visual stimuli preceding the target with short or 0 ISI. Another implication of a delay in the entire whole-brain signal could be that, not only N170, but also other visual evoked potentials, such as P1, N2 and N400 etc., could also be delayed, as long as there is a forward stimulus. Thus, greater caution should be taken when pinpointing a cognitive process to a specific time window when short ISI is used, especially N400 studies (e.g. Kutas & Hillyard, 1980), in which normaly the stimuli of sentences are presented word-by-word in sucession with very short or zero ISI.

### RP is a delayed version of N170

In order to determine if RP is a delayed version of the N170, we measured RP in both the 0-gap condition (with one forward mask, analogous to Dien et al., 2013) and the RSS condition. The former allowed us to determine if RP was really a delayed response, for the 0-gap condition was the end of the continumn of the gap conditions. The latter condition, employing RSS, confirmed that the delayed N1 was same as the typical RP, as the topographical similarity result showed (Figure 7, 8 and Supplementary Figure 3,4,5). Therefore, the results from both conditions have provide the proper evidence for determining whether RP is a delayed version of the N170.

There are a number of implications for an ambiguity between N170 and RP components. Differently labelled components with different latencies are often assumed to reflect different levels of processing. This can be seen in one review that has included RP and N170: Due to latency differences, Dien et al. (2009) relate RP and N170 to different stages in word processing. However, our results now provide us with greater reason to question interpretations like this: Perhaps the same stage of processing was reflected by both components, only with a change in latency. Another implication of RP-N170 ambiguity is that the semantic-processing often associated with RP may prove to be more important to the N170 than is realised in that literature. Most N170 studies focus on the unique visual attributes of a stimulus-category that are often associated with experts. Nonetheless, name-object associations have been found to significantly modulate N170 amplitudes (Heisz & Shedden, 2009); consistent with the idea that the boundary between RP and N170 processing is quite vague. Nonetheless, none of this would be surprising on a physiological level, since we already know that single-cell responses in human IT, believed to underlie much of N170, are sensitive to both visual and semantic attributes (Quiroga, Reddy, Kreiman, Koch, & Fried, 2005). Finally, RP is often described as being more selective for differences in object category, especially familiarity (Martín-Loeches, 2007). While this was mostly conjecture in the past, we have recently shown that, in some conditions, the use of a forward mask can amplify N1-amplitude differences between stimulus categories (Luo et al., 2019). If N170 and RP are not as different as previously thought, then the advantage of using forward-masking as a tool to enhance differential responses should be explored in a wider range studies, especially those in the area of expert object recognition.

This ERP/N170 paper is unusual in that, instead of focusing on stimulus selectivity, we focus on waveform morphology: component latency, width, and topographical distribution. Our parametric manipulation of mask-to-target SOA (gap) allowed us to precisely measure a gradual, continuous change in these morphological features between the opposite ends of what are considered to be ‘N170’ and ‘RP’ conditions. And while the systematic transition between ‘N170’ and ‘RP’ is consistent with the idea that the RP is a delayed version of the N170, form does not necessarily equal function. Nonetheless, we remain optimistic, based on two results. First, our correlation between N1 latency and reaction time suggests that delays in N1 are delays in information processing. Second, we show in another study that the RP and N170 share the same stimulus selectivity in 3 different types of stimulus-category contrasts (Luo et al., 2019). This is an important first step. Ideally, the time-course (and channel distribution) of information processing should be inferred from the data rather than simply assumed to occur at the latencies of pre-defined components (Rousselet et al., 2010; Rousselet & Pernet, 2011);. By taking such an approach, one can determine if ERP sensitivity to stimulus information truly occurs where we expect it to: near the N1 or RP component, depending on the presence of a forward mask and its temporal proximity to the target stimulus. However, that would require a different, more ambitious experimental design, requiring many more trials, and using meaningful stimulus variations that we expect the visual brain to be sensitive to.

One can argue that neither the RSS paradigm nor forward-masking are essential for obtaining RP, they are just tools to amplify RP more effectively (Martín-Loeches, 2007). Let’s take a closer look into the RP studies that didn’t use forward masks. First, in the first RP study by Rudell (1991), the RP with 250 ms latency was a difference wave derived from the subtraction between the target condition presenting one target image on each trial and the control condition displaying only a meaningless image on each trial. This genuine 250 ms RP was, obviously, different from the RP delayed by the forward mask(s). Second, Marí-Beffa, Valdés, Cullen, Catena, & Houghton (2005) didn’t use forward masks and yet claimed they were measuring RP. However, the latency of their RP, about 200 ms, was in the range of N170 latency. Last but not least, the RP observed by Proverbio & Riva (2009), who didn’t use forward masks as well, was in fact the second negative defection, unlike the other RP that was the first. In short, the RPs that were measured without forward masks were either indistinguishable from N170 or distinguishable from the RP measured with forward masks. Since the RP reported by Rudell (see Rudell, 1991 to 2010) was actually a positive wave instead of N1, was that RP also a delayed brain signal? The answer is yes, as long as forward masks were used. Besides, the reason why that RP was a positive wave might be due to the bipolar recording Rudell used.

### Conclusion

In summary, systematic properties of our unique ‘gap-metric’ functions and our time-courses of topographical similarity, all suggest that shorter ISI leads to longer N170 latency, in other words, the delayed N1 should still be recognized as N170. The plausible interpretation of the delay could be the refractory period of the visual system. The implication of the relationship between ISI and the N170 latency also applies for other visual ERPs, that is if a short ISI is used, the component delay should be taken into account when interpreting the time course of the measured effect. Our study also suggests an effective tool to estimate the N170 latency when the peak is hard to identify (i.e., the topographical similarity measurement with dot-product)

Taken together with the results of the gap conditions, the RSS condition further confirmed that the RP waveform is a delayed version of the N170. This implies potential benefits from greater cross-talk between RP and N170 literatures, most especially in how these literatures emphasise different types of information-processing, and in the potential use of forward masks as a tool for enhancing stimulus selectivity of the N170 (Luo et al., 2019). In general, our findings suggest that greater caution should be taken when classifying ERP components based on morphology, and that the effect of specific experimental designs (e.g., ISI, forward masks) on the morphology of these components should be considered when comparing components. But given the flexible nature of component morphology we demonstrate in this study, perhaps morphology-based classification should be replaced or at least validated by data-driven approachs to ERP analysis (e.g., Rousselet et al., 2010; Rousselet & Pernet, 2011).

## Acknowledgments

National Natural Science Foundation of China (NSFC) to Carl M. Gaspar (grant No.: 31371132). This study was funded by the National Natural Science Foundation of China (NSFC) (grant No.: 31400974), Zhejiang Provincial Natural Science Foundation of China (ZJNSF) (grant No.: LY18C090010), and Zhejiang Provincial Dominant and Characteristic Disciplines Cultivation project of College of Education of Hangzhou Normal University (grant No.:9JYXK023) to Ye Zhang.

## Notes

### Competing Interest Statement

The authors have declared no competing interest.

### Summary of Updates

exponential function updated

