## supplemental figures for "Nudging the N170 forward with prior stimulation – Bridging the gap between N170 and recognition potential"

### Supplementary Figures

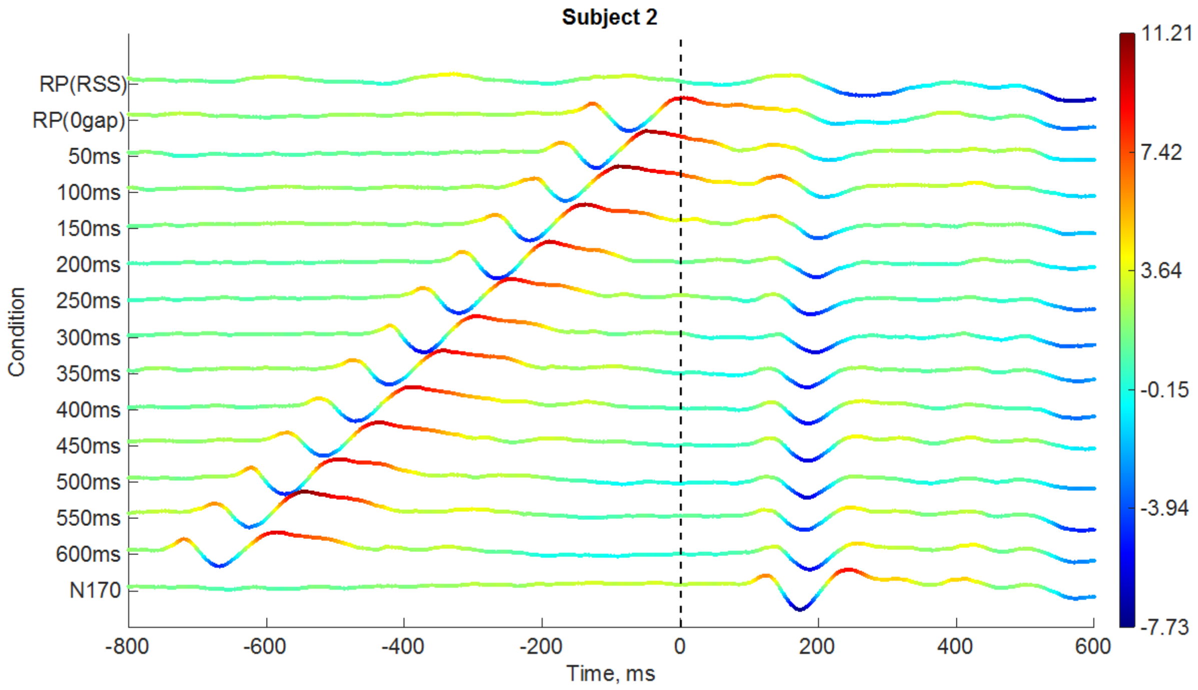


(A)

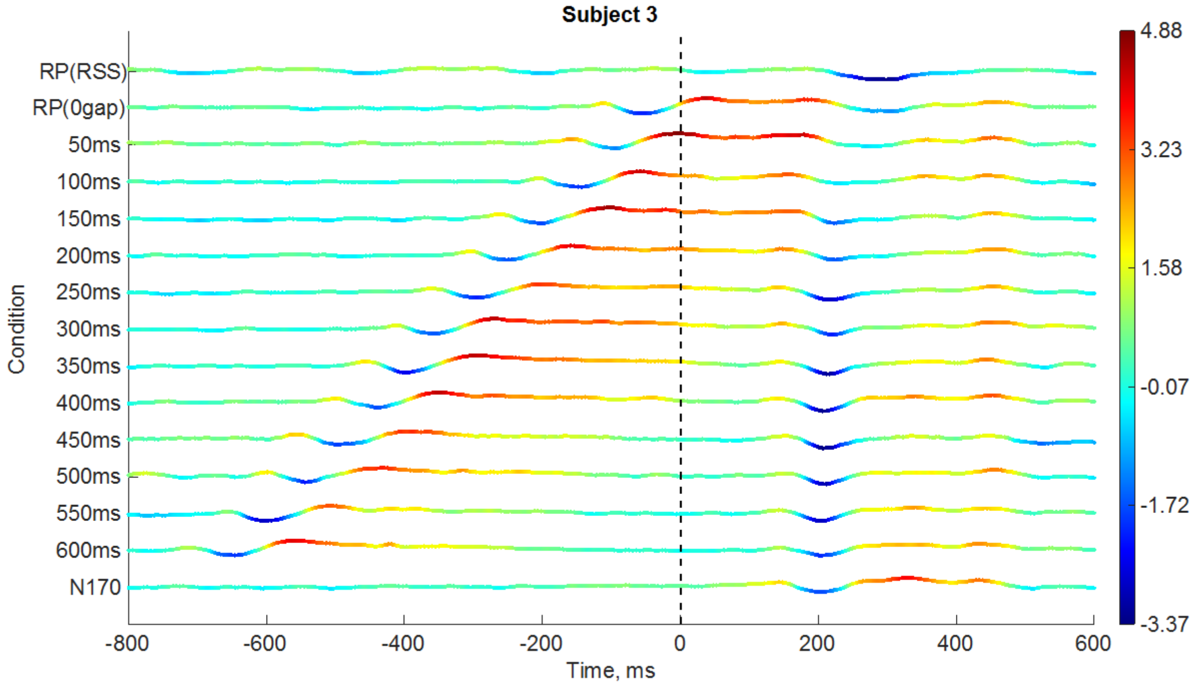
Subject 3
RP(RSS)
RP(0gap)
50ms
100ms
150ms
200ms
250ms
300ms
350ms
400ms
450ms
500ms
550ms
600ms
N170
Condition
-800
-600
-400
-200
0
200
400
600
Time, ms
4.88
3.23
1.58
-0.07
-1.72
-3.37

(B)

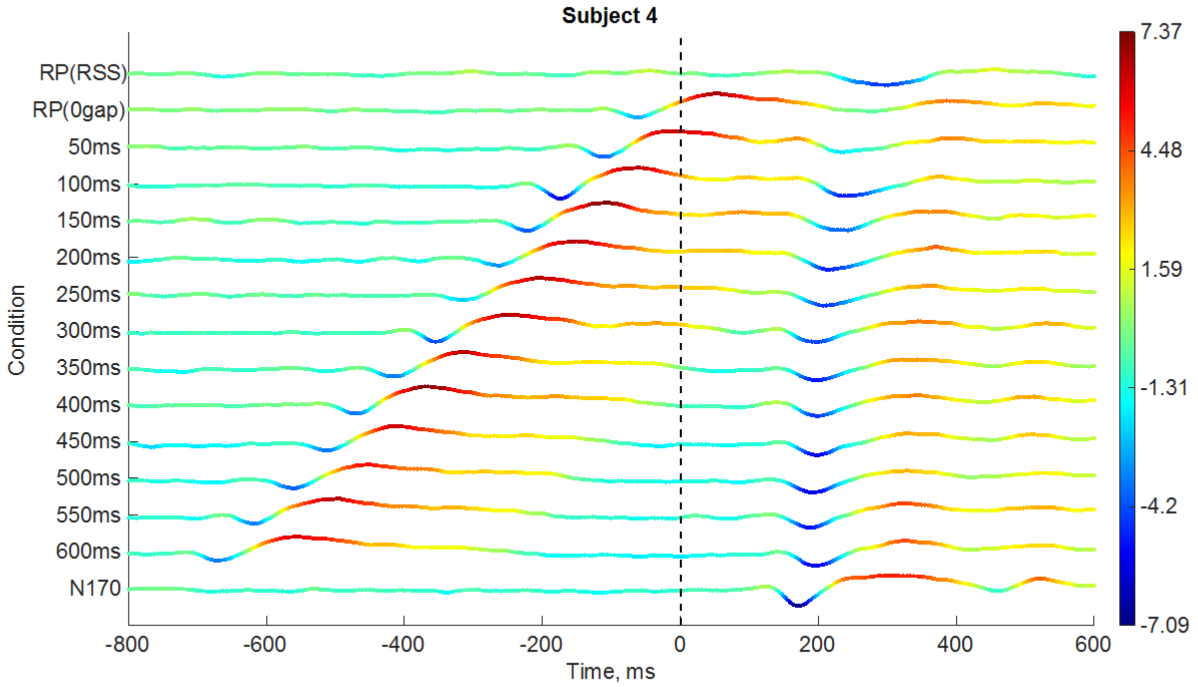
Subject 4
RP(RSS)
RP(0gap)
50ms
100ms
150ms
200ms
250ms
300ms
350ms
400ms
450ms
500ms
550ms
600ms
N170
Condition
-800
-600
-400
-200
0
200
400
600
Time, ms
7.37
4.48
1.59
-1.31
-4.2
-7.09

(C)

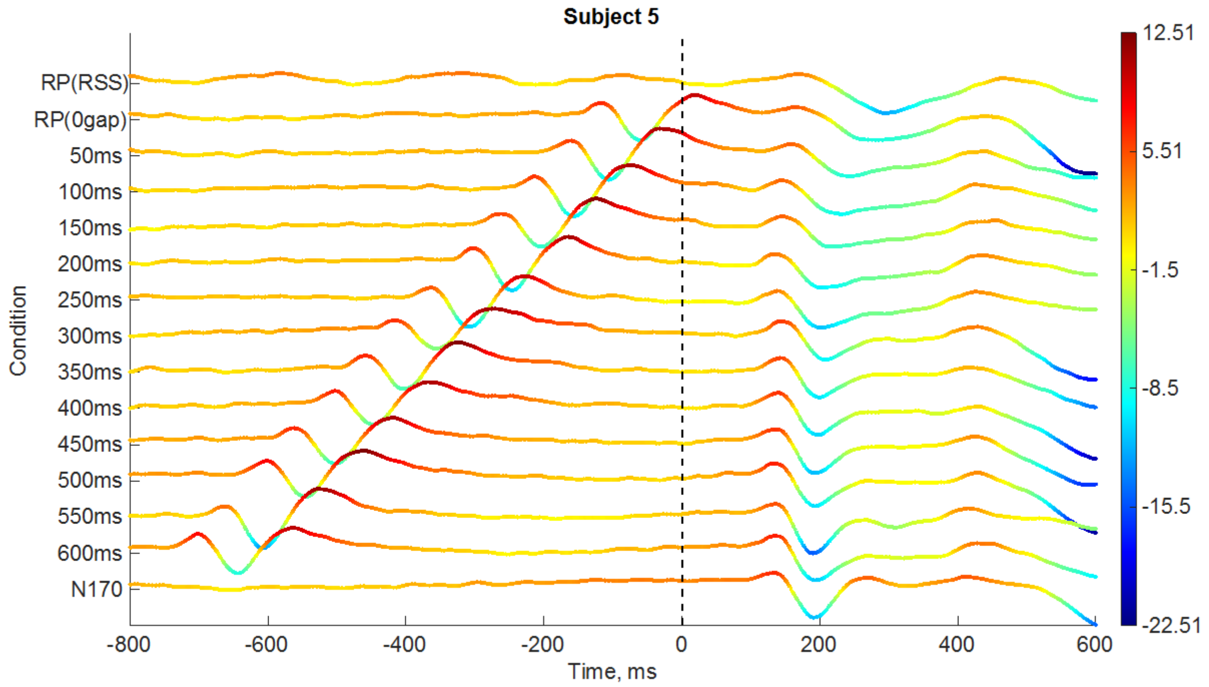
Subject 5
RP(RSS)
RP(0gap)
50ms
100ms
150ms
200ms
250ms
300ms
350ms
400ms
450ms
500ms
550ms
600ms
N170
Condition
-800
-600
-400
-200
0
200
400
600
Time, ms
12.51
5.51
-1.5
-8.5
-15.5
-22.51

(D)

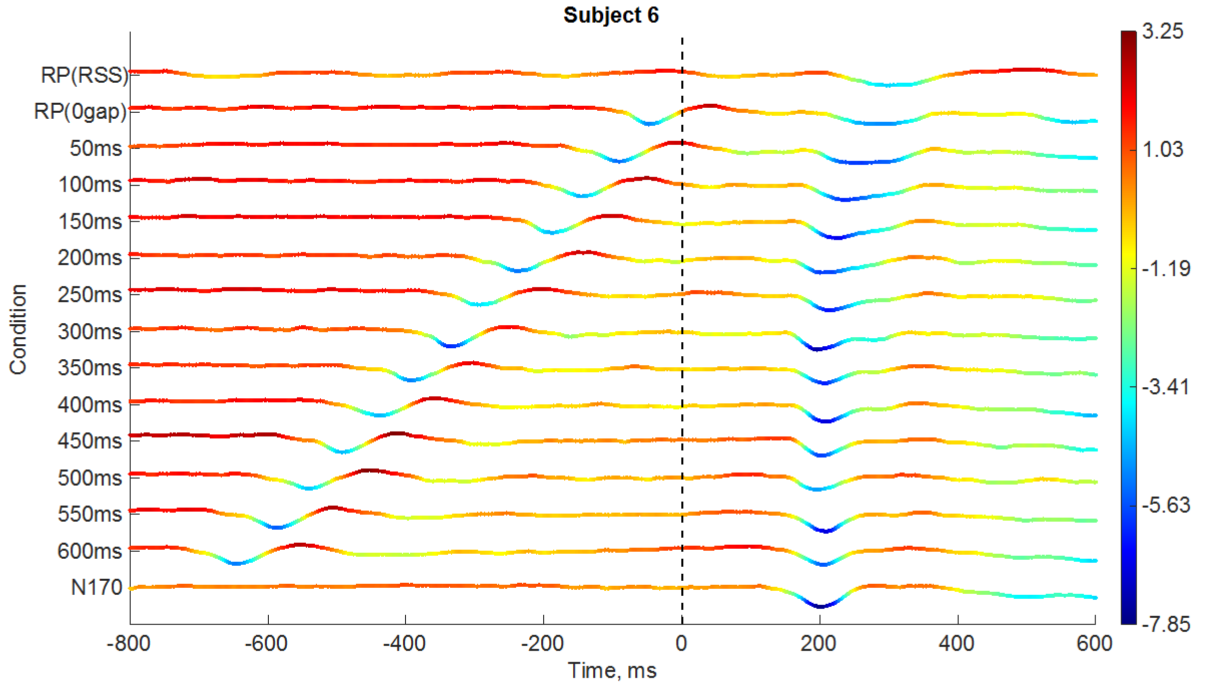
Subject 6
RP(RSS)
RP(0gap)
50ms
100ms
150ms
200ms
250ms
300ms
350ms
400ms
450ms
500ms
550ms
600ms
N170
Condition
-800
-600
-400
-200
0
200
400
600
Time, ms
3.25
1.03
-1.19
-3.41
-5.63
-7.85

(E)

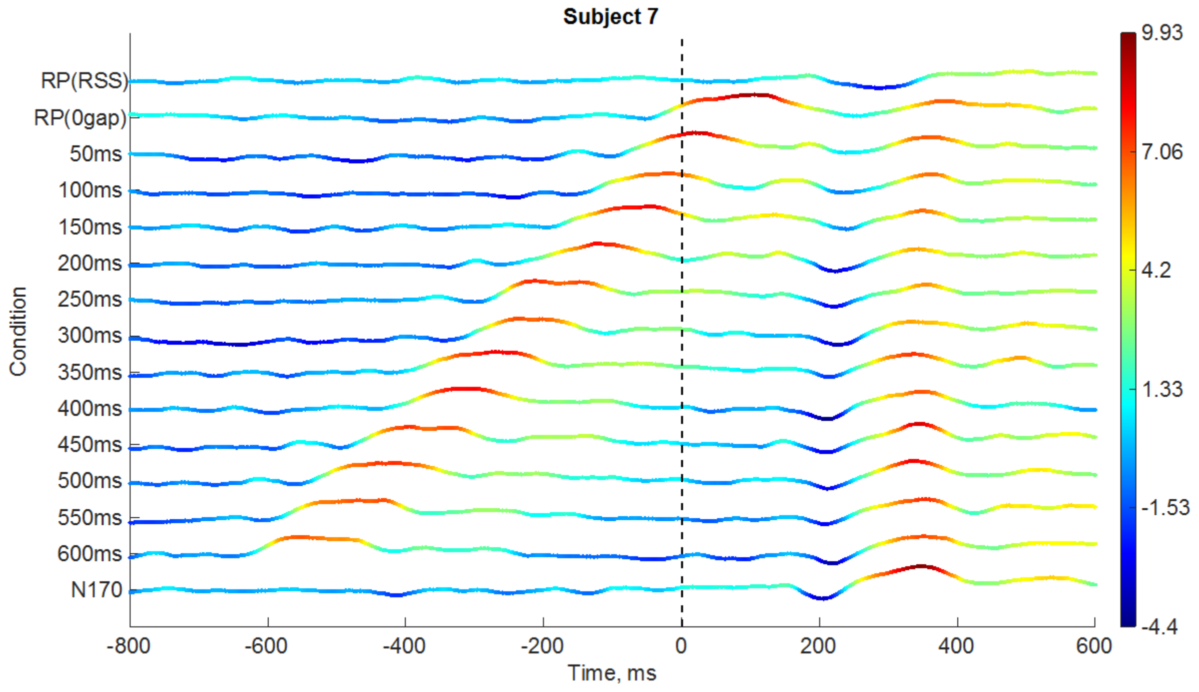


(F)

**Supplementary Figure 1.** ERP stack plots for subjects S2 to S7. Figures for S1 is in the main text. Mean ERP are stacked vertically across conditions and voltage is colour-coded, with a corresponding colour-bar to the right.

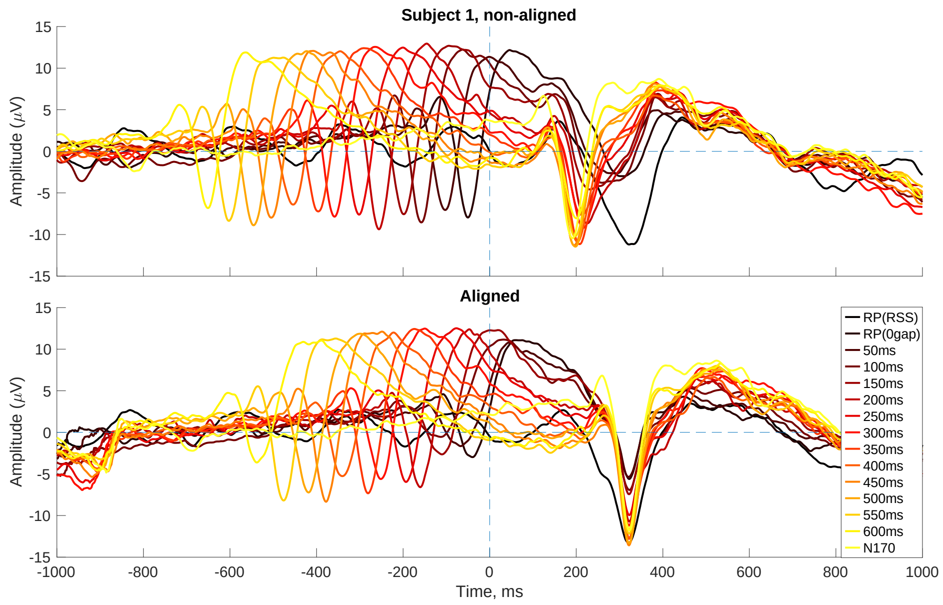


(A)

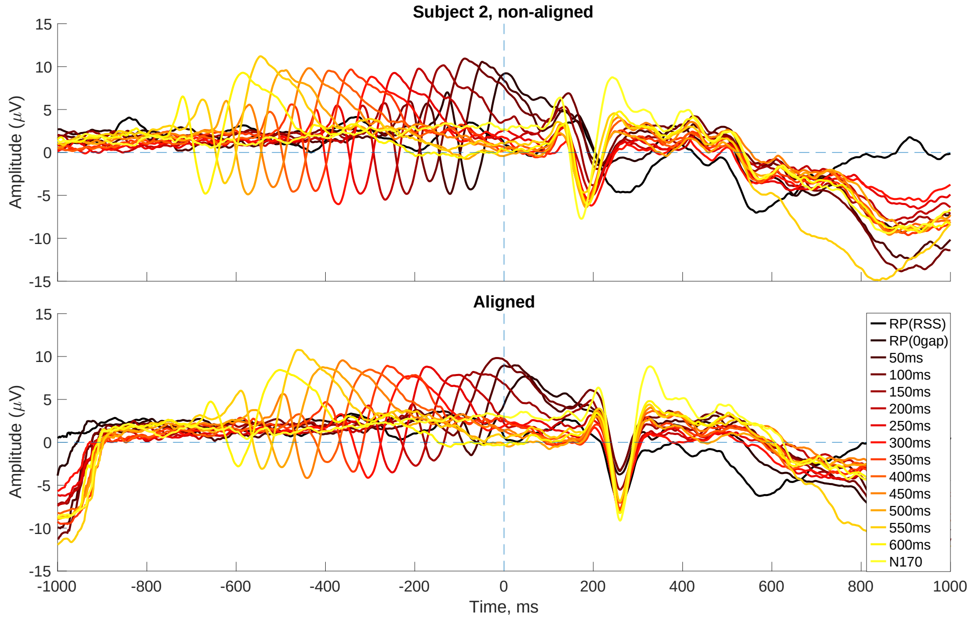
Subject 2, non-aligned
Aligned
Amplitude (μV)
Time, ms
RP(RSS)
RP(0gap)
50ms
100ms
150ms
200ms
250ms
300ms
350ms
400ms
450ms
500ms
550ms
600ms
N170

(B)

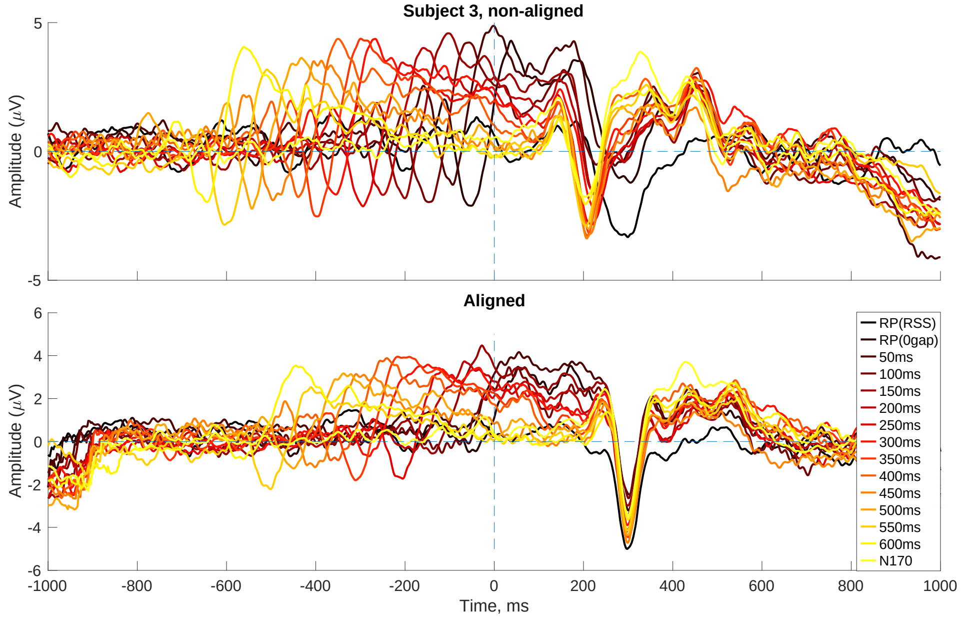
Subject 3, non-aligned
Aligned
Amplitude (μV)
Time, ms
RP(RSS)
RP(0gap)
50ms
100ms
150ms
200ms
250ms
300ms
350ms
400ms
450ms
500ms
550ms
600ms
N170

(C)

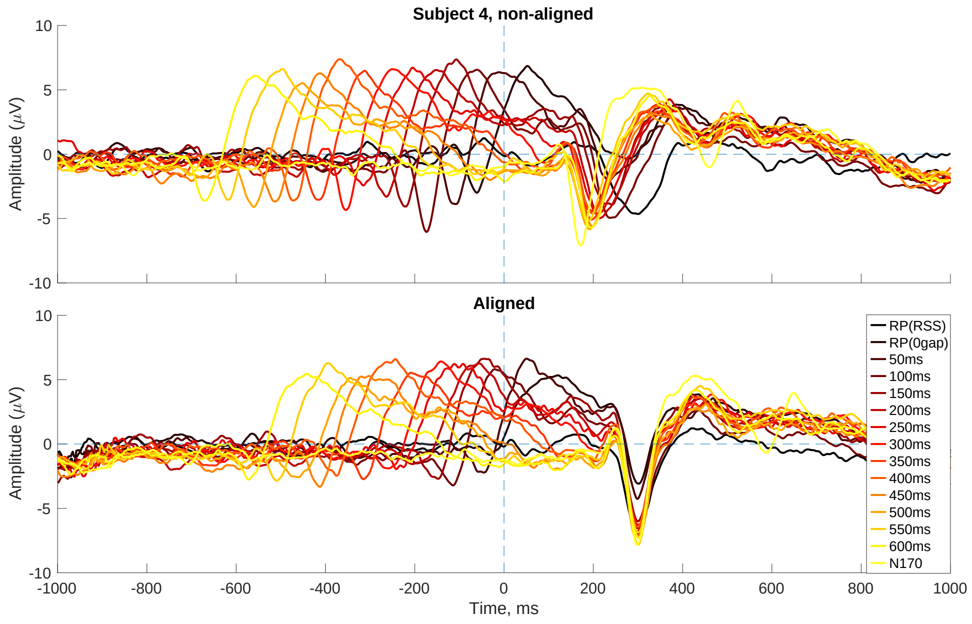
Subject 4, non-aligned
Aligned
Amplitude (μV)
Time, ms
RP(RSS)
RP(0gap)
50ms
100ms
150ms
200ms
250ms
300ms
350ms
400ms
450ms
500ms
550ms
600ms
N170

(D)

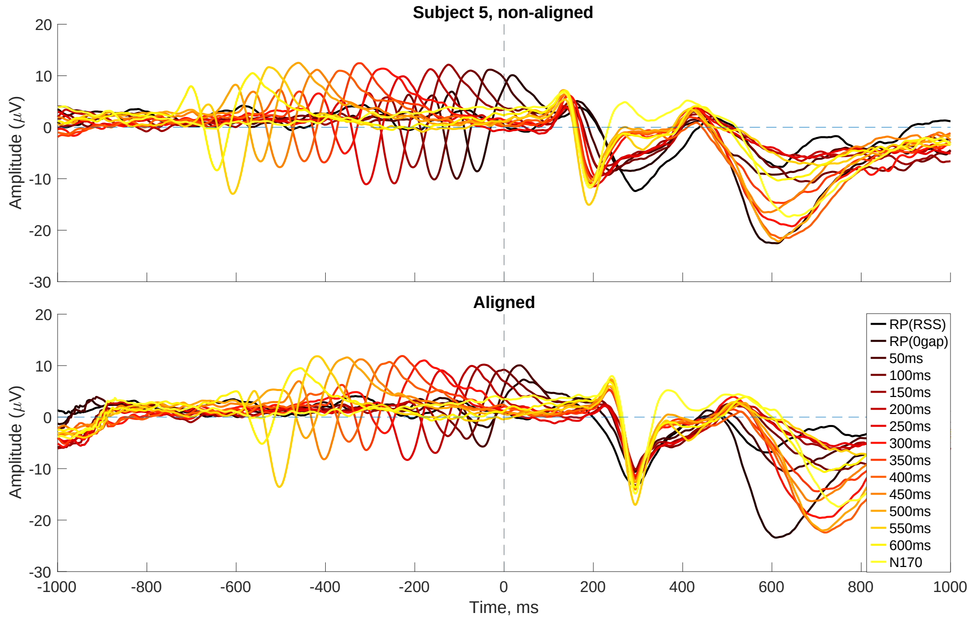
Subject 5, non-aligned
Aligned
Amplitude (μV)
Time, ms
RP(RSS)
RP(0gap)
50ms
100ms
150ms
200ms
250ms
300ms
350ms
400ms
450ms
500ms
550ms
600ms
N170

(E)

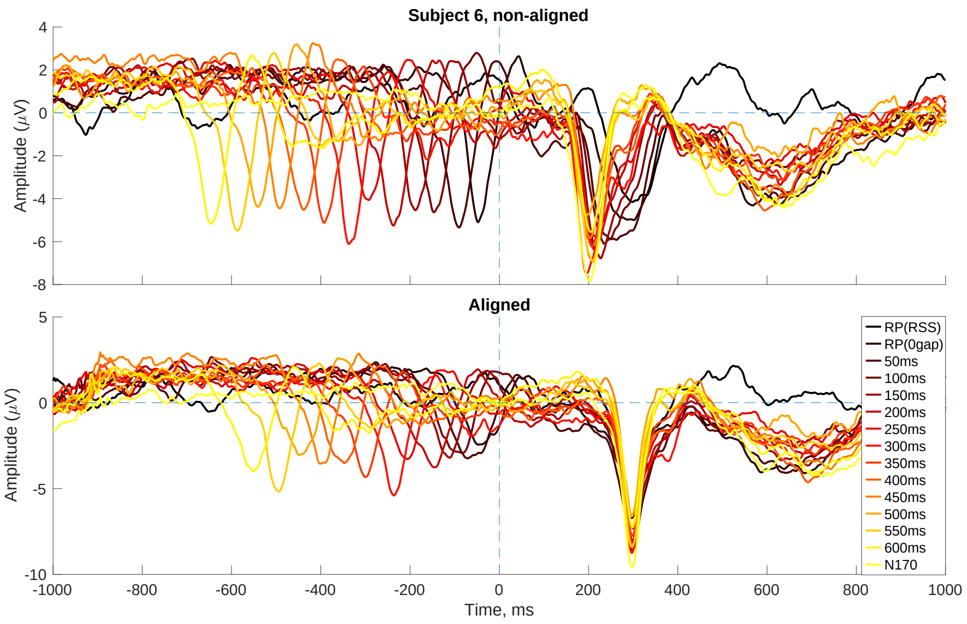


(F)

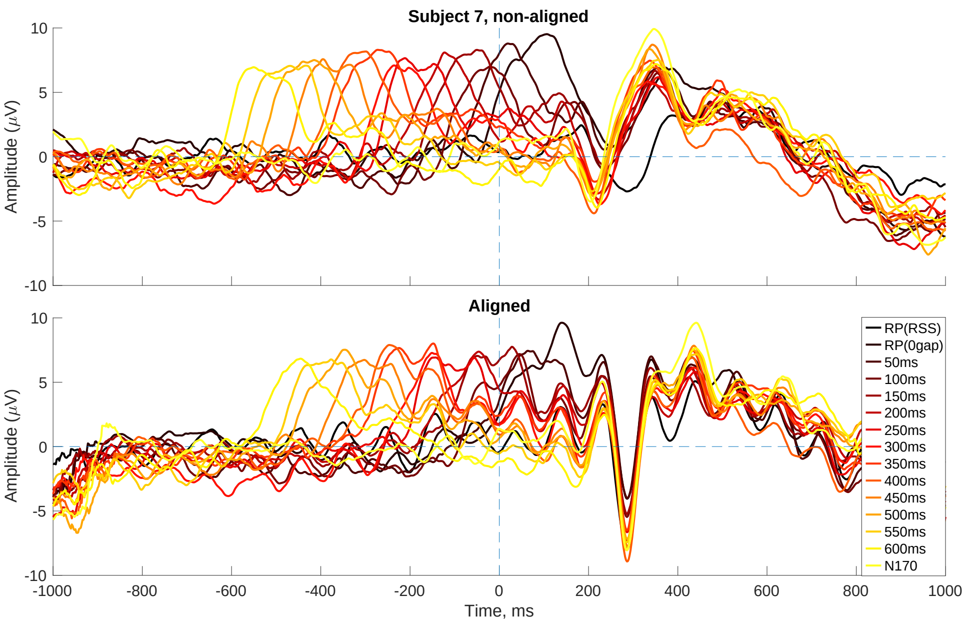


(G)

**Supplementary Figure 2.** Average N1 after aligning single-trial peaks for subjects S1 to S7. For each subplot, the top panel is the original ERPs (15 conditions are plotted on the same y axis), the bottom panel is the ERPs that the single-trial peaks for each condition were aligned to a same time point.

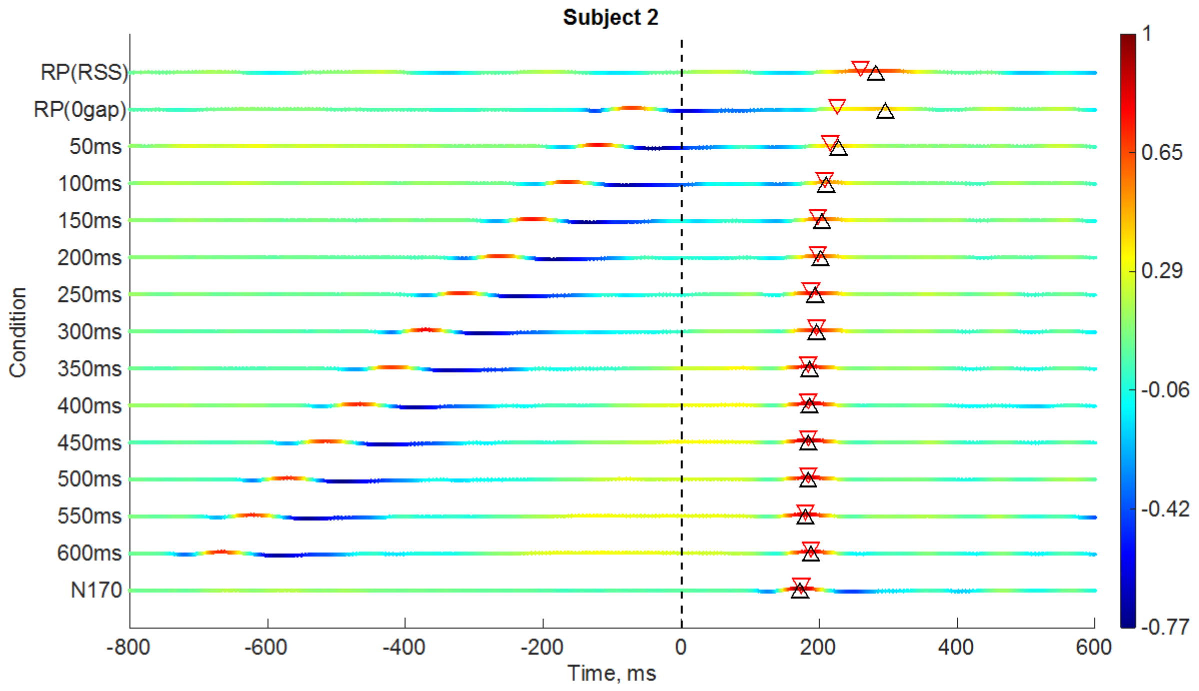


**(A)**

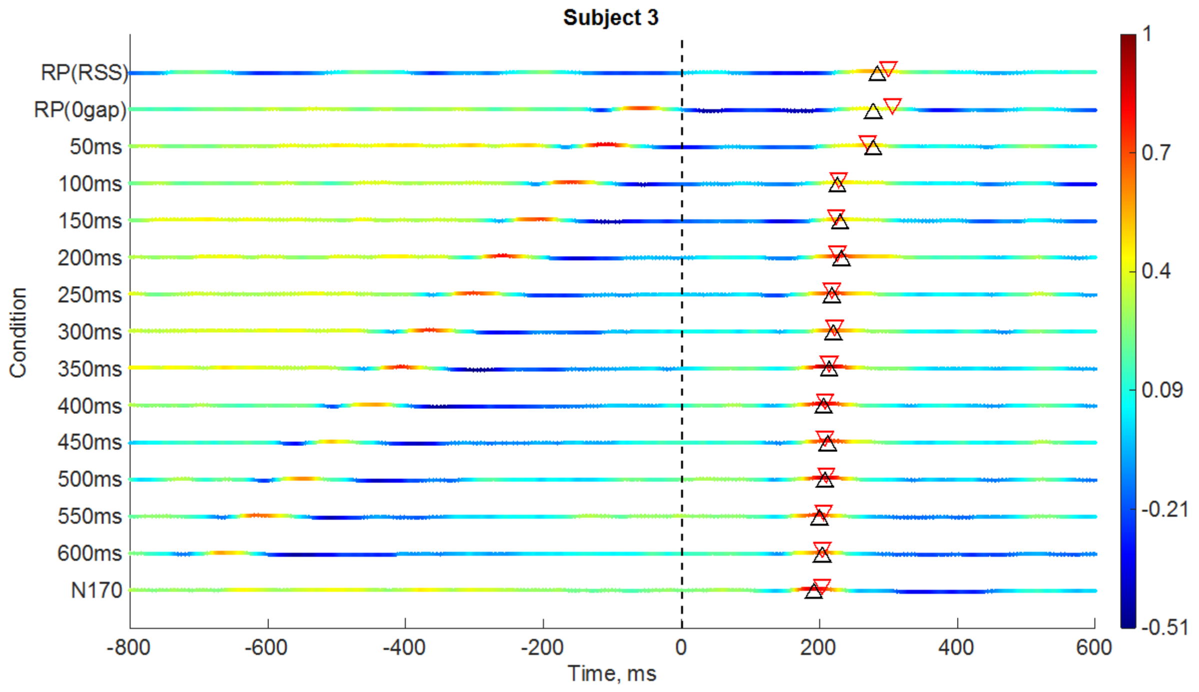

(B)

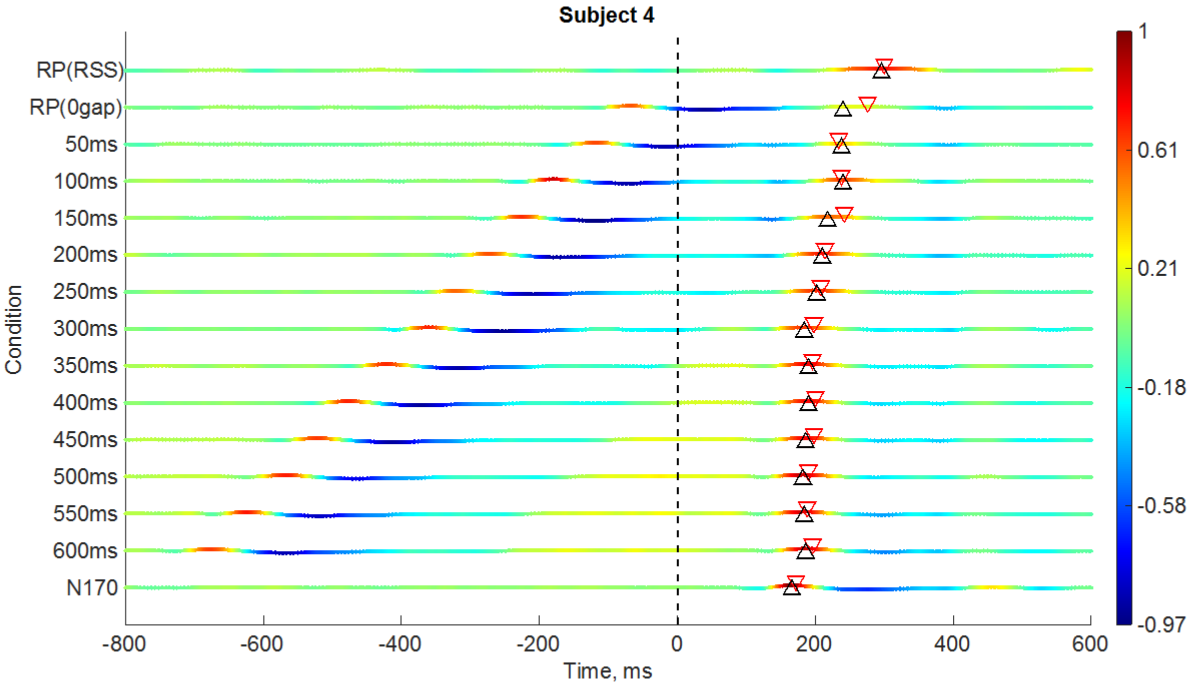
Subject 4
RP(RSS)
RP(0gap)
50ms
100ms
150ms
200ms
250ms
300ms
350ms
400ms
450ms
500ms
550ms
600ms
N170
Condition
-800
-600
-400
-200
0
200
400
600
Time, ms
1
0.61
0.21
-0.18
-0.58
-0.97

(C)

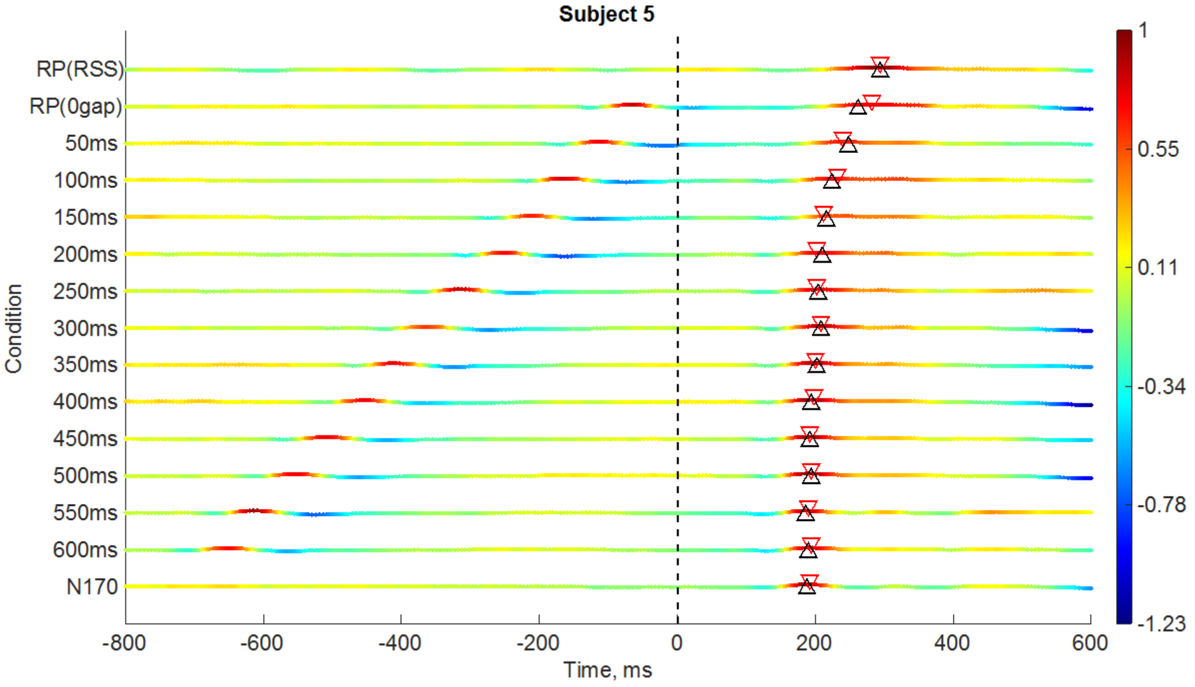
Subject 5
RP(RSS)
RP(0gap)
50ms
100ms
150ms
200ms
250ms
300ms
350ms
400ms
450ms
500ms
550ms
600ms
N170
Condition
-800
-600
-400
-200
0
200
400
600
Time, ms
1
0.55
0.11
-0.34
-0.78
-1.23

(D)

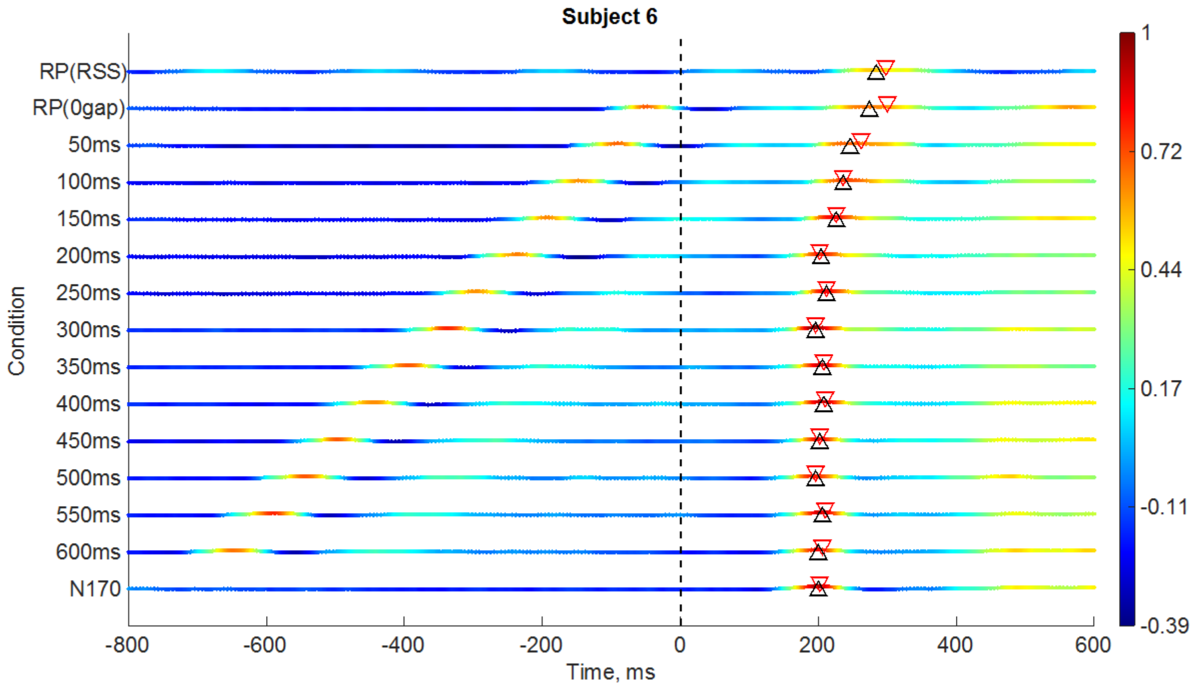


**(E)**

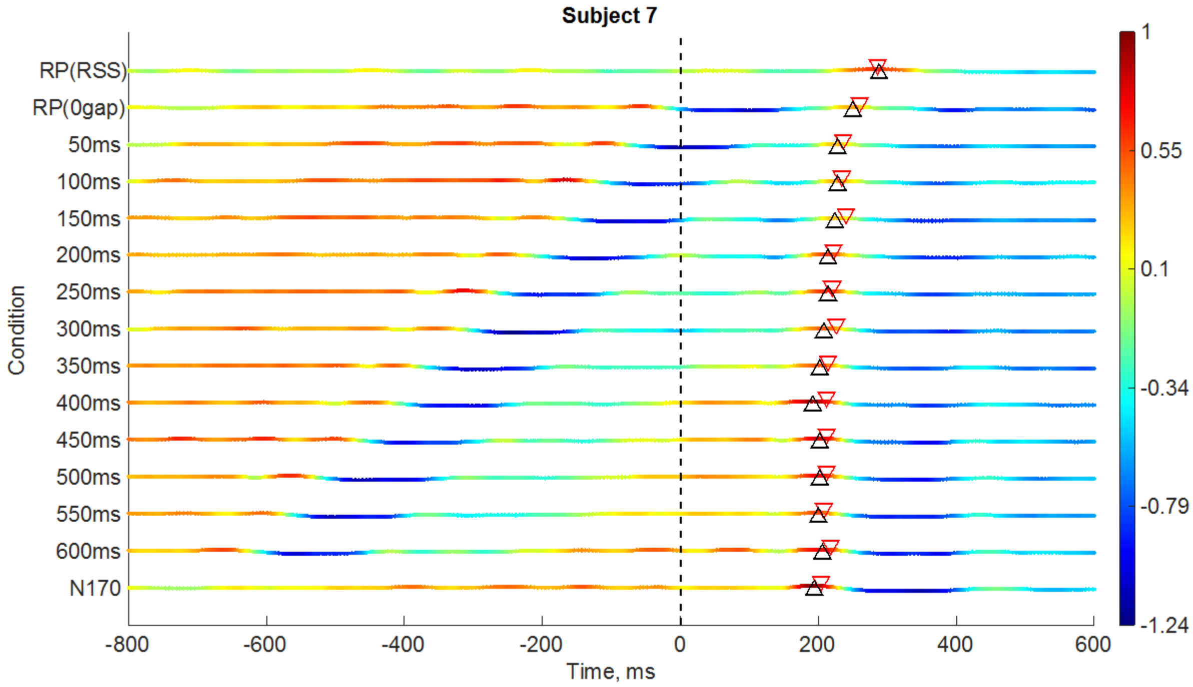


**(F)**

**Supplementary Figure 3.** Time-course of inner product between N170 topographical distribution (at N1 latency, as shown in the top row of Figure 7 of main text), and topographies at every other time-frame, and condition (figure of S1 is in the main text). To improve visualisation, each inner-product time-course is normalised by its maximum inner-product. In each condition, a black triangle (pointing up) shows the latency of the N1 for the mean ERP in that condition; and a red triangle (pointing down) shows the latency of the topography in that condition which has the highest inner-product with the N170 topography (at N1), within 0 to 400 ms. According to our hypothesis, black and red triangles should roughly match up.

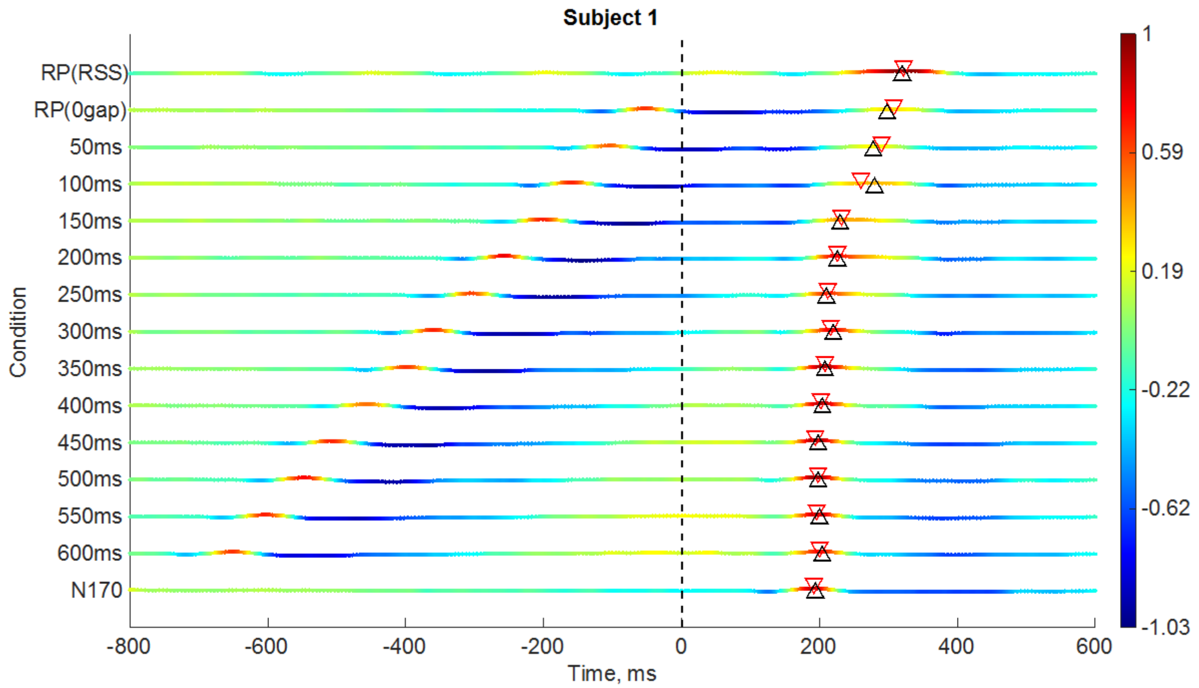


**(A)**

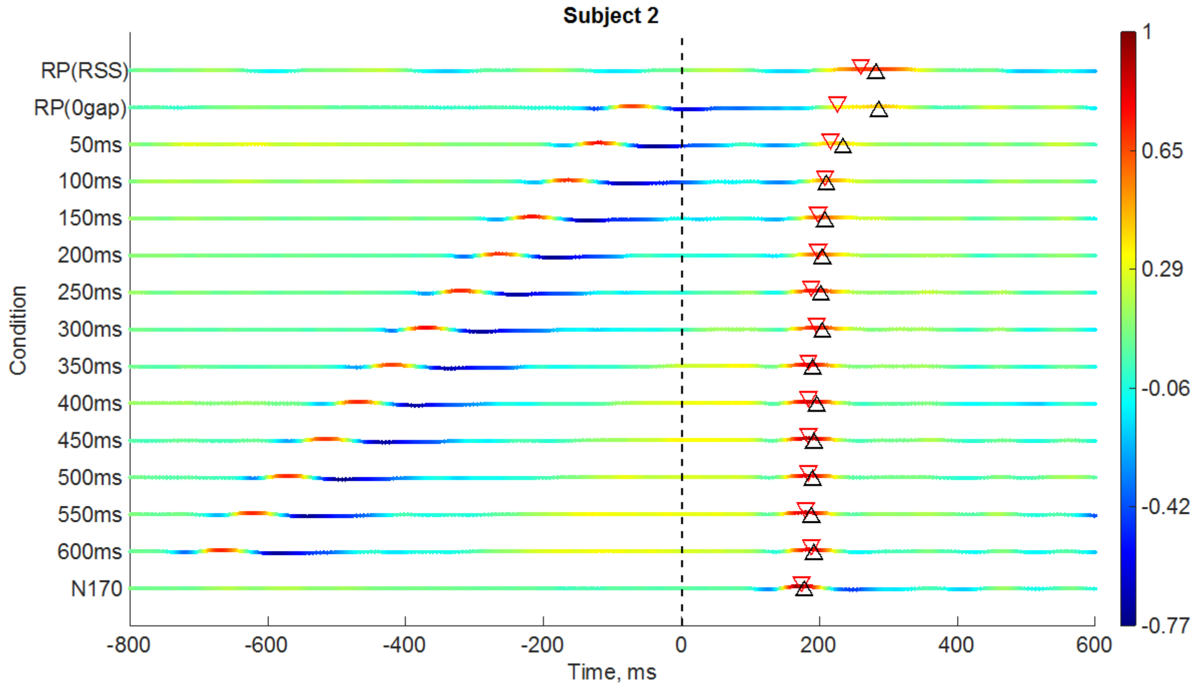


**(B)**

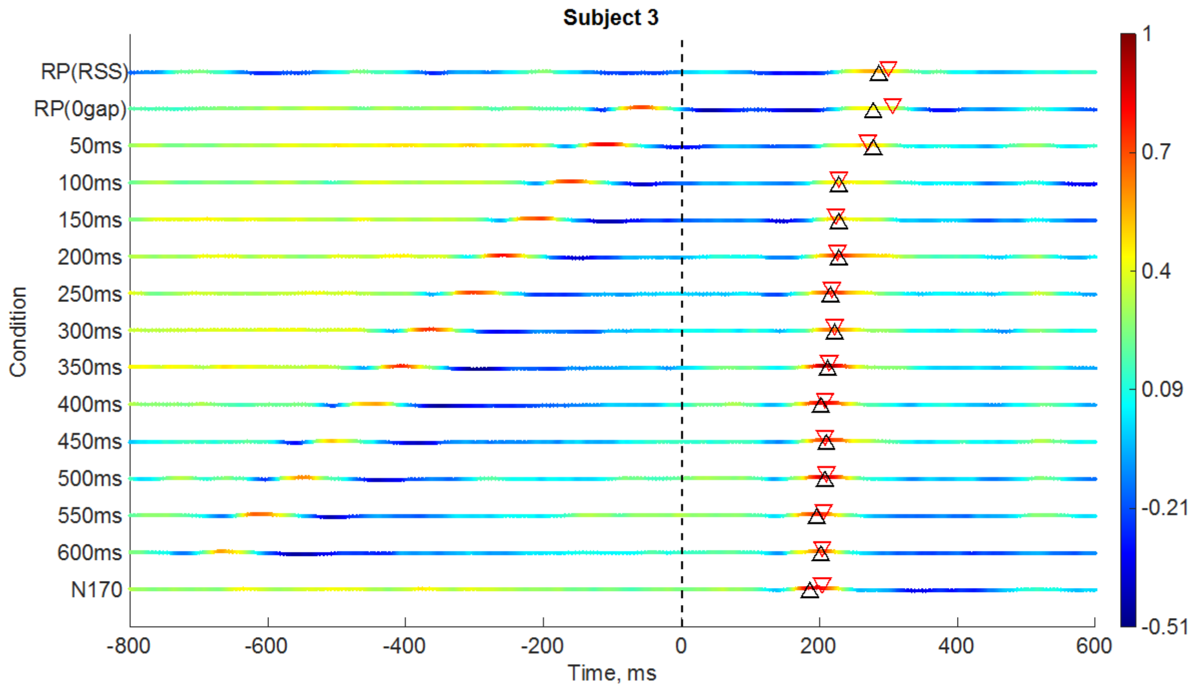
Subject 3
RP(RSS)
RP(0gap)
50ms
100ms
150ms
200ms
250ms
300ms
350ms
400ms
450ms
500ms
550ms
600ms
N170
Condition
-800
-600
-400
-200
0
200
400
600
Time, ms
1
0.7
0.4
0.09
-0.21
-0.51

**(C)**

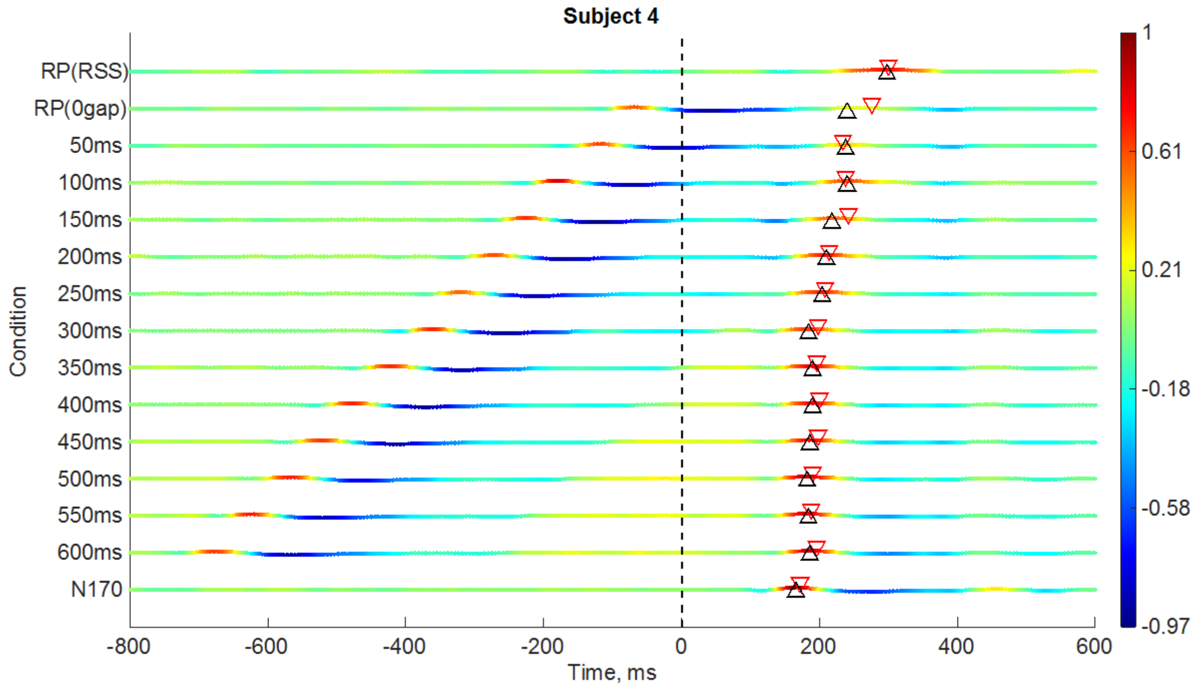
Subject 4
RP(RSS)
RP(0gap)
50ms
100ms
150ms
200ms
250ms
300ms
350ms
400ms
450ms
500ms
550ms
600ms
N170
Condition
-800
-600
-400
-200
0
200
400
600
Time, ms
1
0.61
0.21
-0.18
-0.58
-0.97

**(D)**

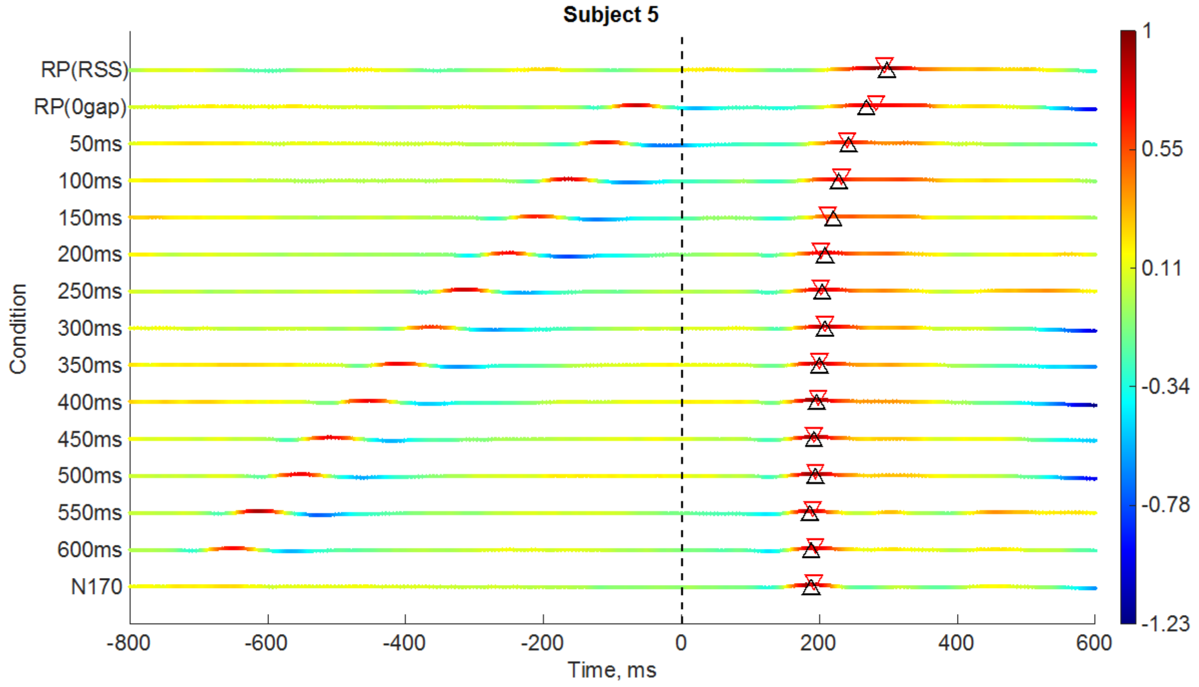
Subject 5
RP(RSS)
RP(0gap)
50ms
100ms
150ms
200ms
250ms
300ms
350ms
400ms
450ms
500ms
550ms
600ms
N170
Condition
-800
-600
-400
-200
0
200
400
600
Time, ms
1
0.55
0.11
-0.34
-0.78
-1.23

**(E)**

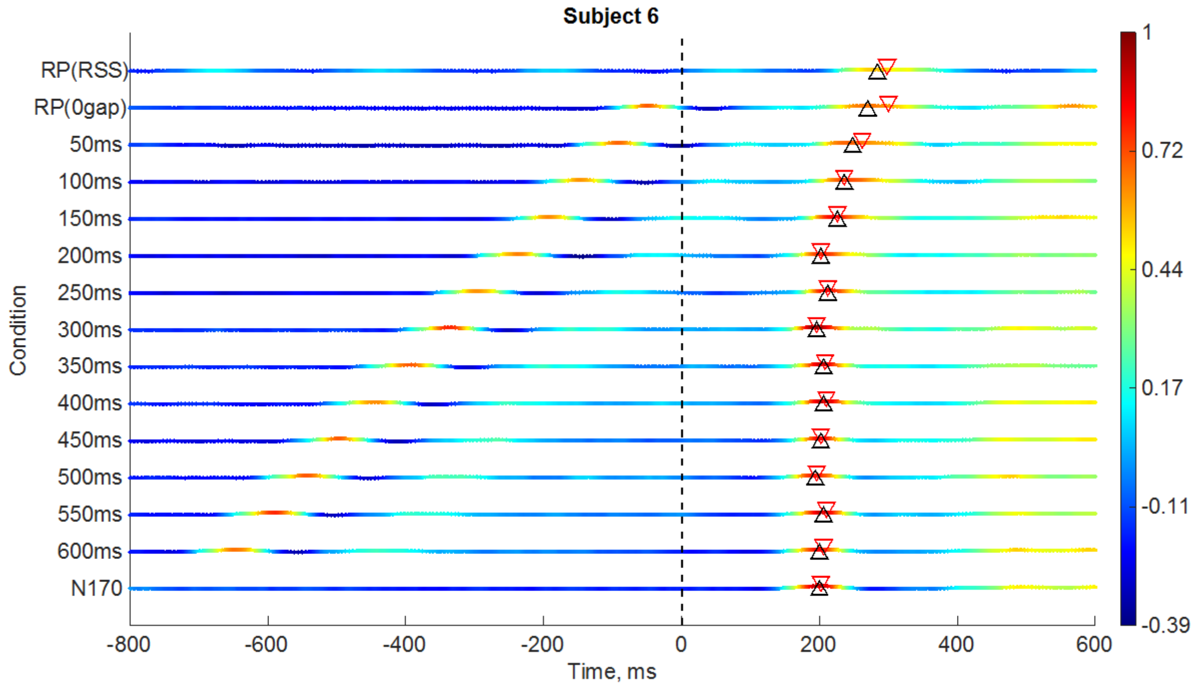
Subject 6
RP(RSS)
RP(0gap)
50ms
100ms
150ms
200ms
250ms
300ms
350ms
400ms
450ms
500ms
550ms
600ms
N170
Condition
-800
-600
-400
-200
0
200
400
600
Time, ms
1
0.72
0.44
0.17
-0.11
-0.39

**(F)**

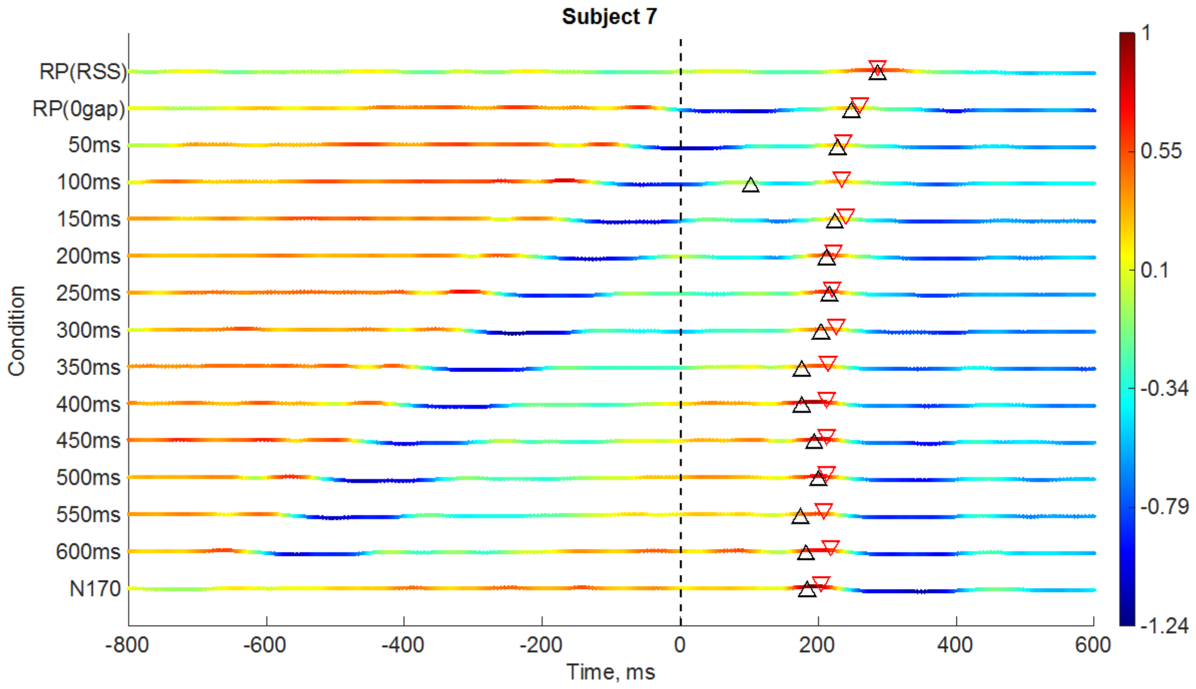


**(G)**

**Supplementary Figure 4.** Time-course of inner product between RP (RSS condition) topographical distribution (at RP latency), and topographies at every other time-frame, and condition. To improve visualisation, each inner-product time-course is normalised by its maximum inner-product. In each condition, a black triangle (pointing up) shows the latency of the N1 for the mean ERP in that condition; and a red triangle (pointing down) shows the latency of the topography in that condition which has the highest inner-product with the RP topography (at N1), within 0 to 400 ms.

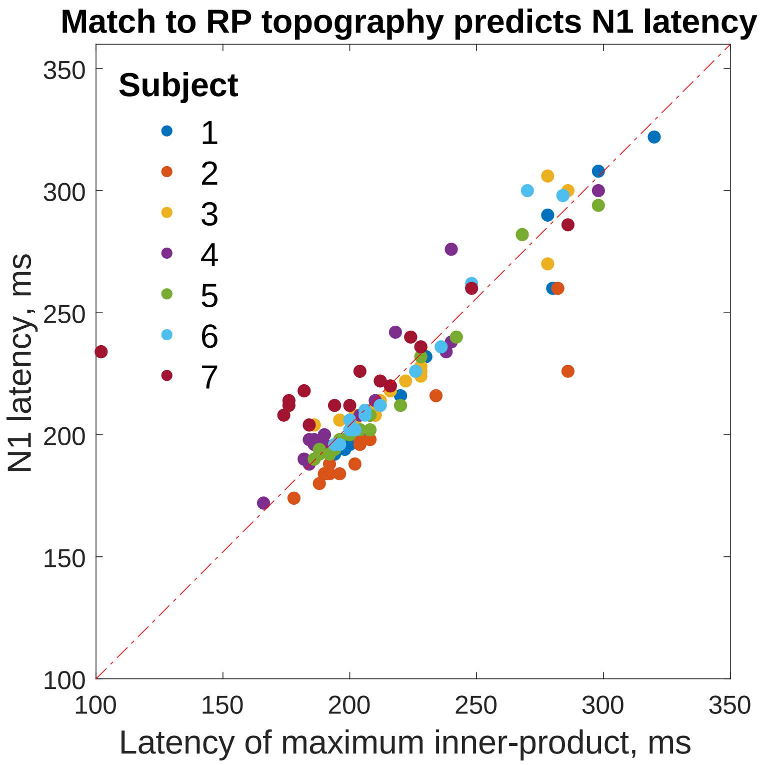


**Supplementary Figure 5.** The latency of maximum inner-product with RP (RSS condition) topography (at N1) for each gap condition, predicts the N1 latency for that condition, for each subject. ($r = 0.8620$, $p = 3.6149e-32$)
